# Heritable CRISPR-Cas9 editing of plant genomes using RNA virus vectors

**DOI:** 10.1101/2022.09.07.507055

**Authors:** Mireia Uranga, Verónica Aragonés, José-Antonio Daròs, Fabio Pasin

**Affiliations:** Instituto de Biología Molecular y Celular de Plantas (IBMCP), Consejo Superior de Investigaciones Científicas – Universitat Politècnica de València, Avenida de los Naranjos s/n, 46022 Valencia, Spain; VIB-UGent Center for Plant Systems Biology, Technologiepark-Zwijnaarde 71, 9052 Zwijnaarde (Gent), Belgium

## Abstract

Viral vectors hold enormous potential for genome editing in plants as transient delivery vehicles of CRISPR-Cas components. Here, we describe a protocol to assemble plant viral vectors for single guide RNA (sgRNA) delivery. The obtained viral constructs are based on compact T-DNA binary vectors of the pLX series and are delivered into Cas9-expressing plants through agroinoculation. This approach allows rapidly assessing sgRNA design for plant genome targeting, as well as the recovery of progeny with heritable mutations at targeted loci.

**For complete details on the use and execution of this protocol, please refer to** Uranga et al.^1^ **and** Aragonés et al.^2^.

**Graphical abstract:** 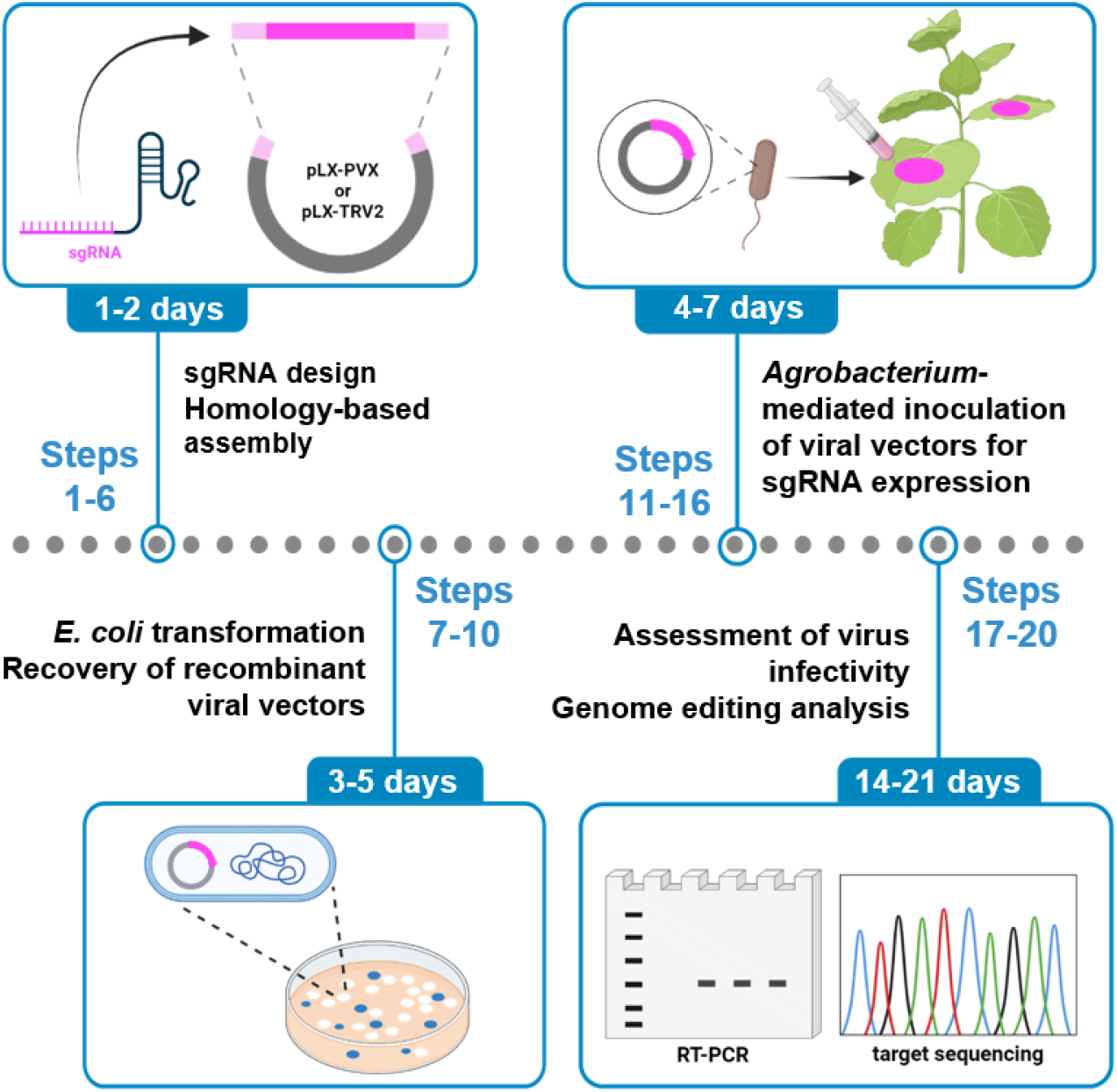

## Before you begin

Protocols and computational tools were reported for CRISPR/Cas genome editing^3–5^. Viral vectors are a promising alternative for the transient delivery of the CRISPR-Cas system in many biological systems including plants, following the so-called virus-induced genome editing (VIGE) approach^6–8^. The systemic spread of the viral infection promotes an accumulation of the editing components within plant tissues. This leads to efficient and fast genome editing, thus providing an ideal screening tool to assess the effectivity and specificity of single guide RNA (sgRNA) design. Several plant RNA virus-based replicons have been successfully used for the delivery of sgRNAs in transgenic plants constitutively expressing the Cas9 nuclease^9–19^. However, each viral vector has its own molecular biology properties and is restricted to a specific host range. Here, we describe the engineering of two viral vectors derived from potato virus X (PVX; genus *Potexvirus*) and tobacco rattle virus (TRV; genus *Tobravirus*) for the delivery of unspaced sgRNAs in the model species *Nicotiana benthamiana* (Figure 1). The presented PVX system consists of a single binary vector pLX-PVX, which includes PVX genomic sequences and a heterologous sub-genomic promoter from bamboo mosaic virus (BaMV) to drive insert expression (Figure 2). The TRV system relies on pLX-TRV1 and pLX-TRV2, two T-DNA vectors with compatible origins for simultaneous agroinoculation of viral genomic components (JoinTRV). pLX-TRV1 provides the replicase function, whereas pLX-TRV2 includes an engineered TRV RNA2 sequence with a heterologous sub-genomic promoter of pea early browning virus (PEBV) to drive insert expression (Figure 2). Both viral systems are based on compact T-DNA binary vectors of the pLX series^20^, which have been successfully used for starting RNA and DNA virus infections by *Agrobacterium*-mediated inoculation (agroinoculation)^21–23^. Recombinant viral replicons with sgRNA constructs are assembled and delivered into Cas9-expressing plants via agroinoculation. Systemic viral infection results in germline genome editing and recovery of edited progeny (Figure 1).

**Figure 1.**
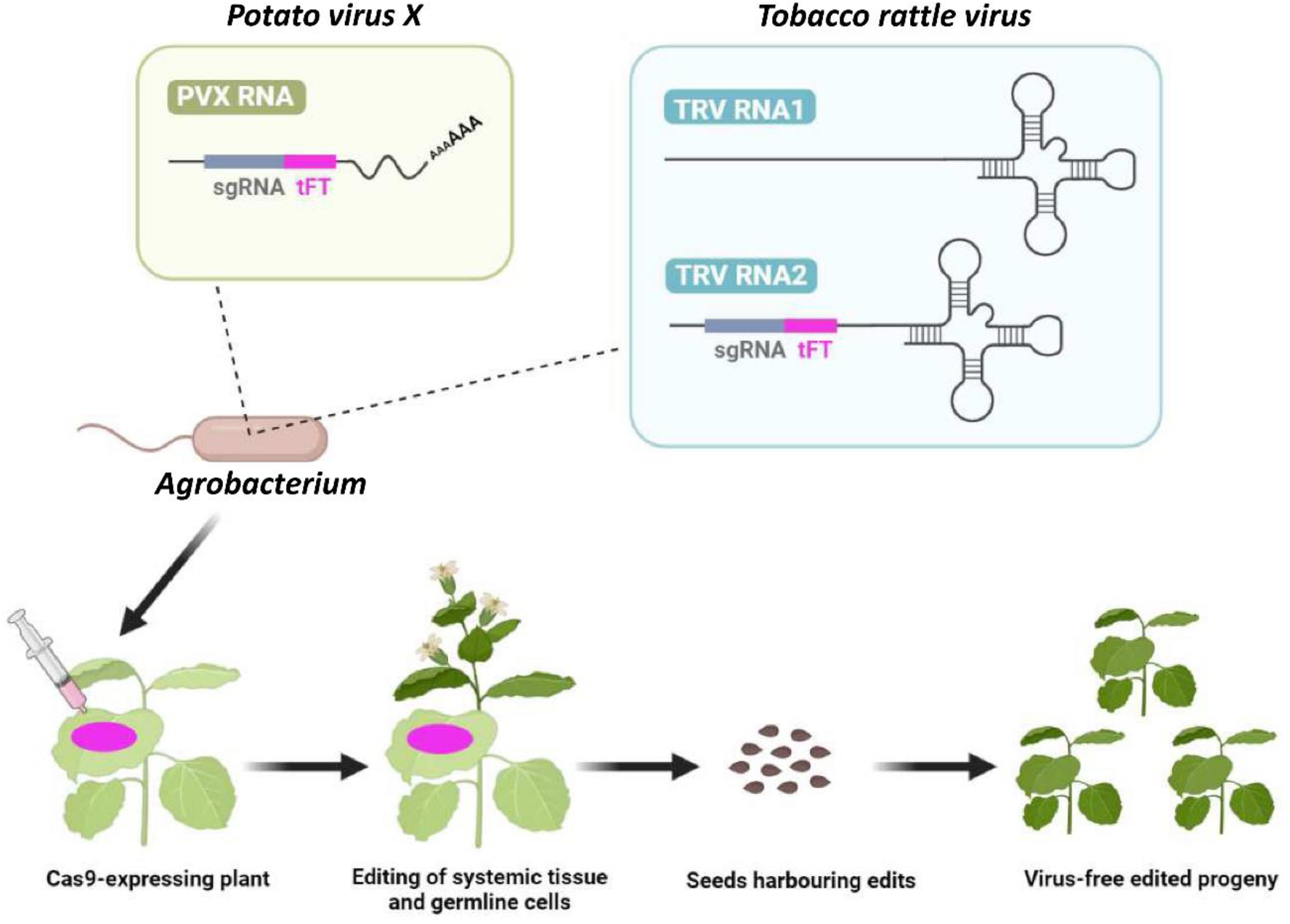
Protocol outline for CRISPR-Cas9-mediated genome editing in plants using viral vectors. Cas9-expressing *N. benthamiana* plants are agroinoculated with an RNA viral vector carrying a single guide RNA (sgRNA) fused to an RNA mobility element from *A. thaliana Flowering locus T* (tFT). Systemic viral infection results in germline genome editing and recovery of edited progeny.

**Figure 2.**
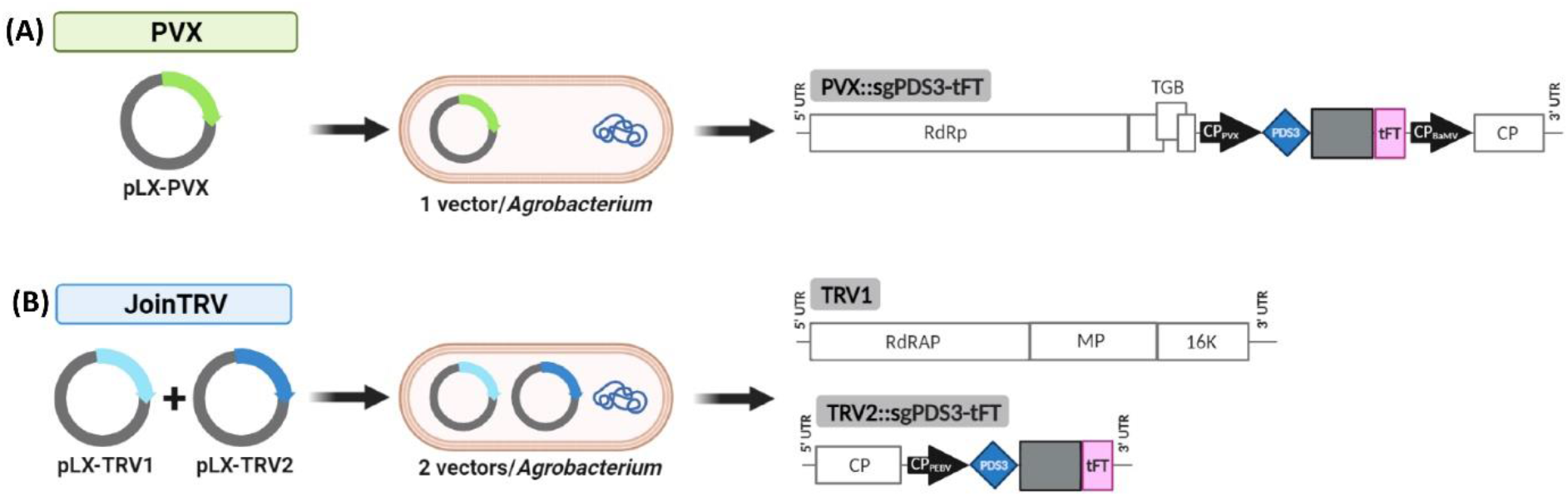
Schematic representation of the viral vectors used. Viral vectors for delivery of sgPDS3-tFT, a single guide RNA (sgRNA) construct that comprises of a protospacer (blue diamond) targeting both *N. benthamiana PDS* homeologs, a conserved Cas9 scaffold (gray box) and a truncated *A. thaliana Flowering locus T* (tFT; a pink box). **(A)** Diagram of the PVX vector system^1^. An *Agrobacterium* strain hosts pLX-PVX, a T-DNA vector with the pBBR1 origin and kanamycin resistance for PVX agroinoculation (see Supporting Information). **(B)** Diagram of the JoinTRV system^2^. An *Agrobacterium* strain simultaneously hosts pLX-TRV1 and pLX-TRV2, two compatible T-DNA vectors for TRV agroinoculation. pLX-TRV1 is a derivative of pLX-Z4, which includes the RK2 origin and gentamicin resistance^20^; pLX-TRV2 is a derivative of pLX-B2, which includes the pBBR1 origin and kanamycin resistance^20^. RdRp, RNA-dependent RNA polymerase; TGB, triple gene block; CP, coat protein; RdRAP, RNA-dependent replication-associated protein; MP, movement protein; 16K, 16 kDa protein. Black arrows show the heterologous bamboo mosaic virus (BaMV) and pea early browning virus (PEBV) coat protein promoters; promoters, sgRNA and tFT, not at scale.

### Biosafety

Bacteria containing recombinant nucleic acid molecules need to be handled and disposed according to institutional regulations. Plant viruses and derived vectors pose potential agronomical and environmental risks; therefore, researchers must comply with governmental and institutional regulations for virus propagation in plants and working with plant infectious agents. Additional institutional regulations must be considered when using transgenic plant lines such as the Cas9-expressing *N. benthamiana* described here.

### Cas9-expressing plants and growth conditions

#### Timing: 4-5 weeks

Start preparing the plant material four to five weeks in advance as follows:

1. Sow *N. benthamiana* Cas9 seeds in a well-watered soil mixture (one part of perlite and two parts of potting substrate) and let them germinate in a growth chamber set at 25°C and long-day conditions (16 h-light/8 h-dark). **CRITICAL:** A variety of Cas nuclease effectors have been reported^6^. This protocol has been validated using a *N. benthamiana* transgenic line that constitutively expresses a human codon-optimized version of *Streptococcus pyogenes* Cas9 fused to a nuclear localization signal (NLS) at the carboxyl terminus. The Cas9 gene is under the control of the cauliflower mosaic virus 35S promoter and *A. tumefaciens nopaline synthase* terminator^1^.
2. Two weeks after sowing, transfer three seedlings per plastic pot filled with the same soil mixture and water them every 2 days to ensure that the soil is humid. ***CRITICAL:*** Avoiding water excess in the tray.
3. Four- to five-week-old plants (i.e. five-leaf stage) are ideal for *Agrobacterium*-mediated inoculation of viral vectors.

***CRITICAL***: The quality and age of the plants are essential factors for observing homogeneous symptoms of systemic viral infection and achieving successful genome editing. Healthy plants are obtained only following good practices in a well-conditioned growth chamber, including a disease- and pest-free environment and careful watering.

***Note:*** Plants can grow at different pace in different places, therefore following a leaf stage can be more appropriate than an age stage. For optimal agroinoculation conditions and time saving, ensure to coordinate the preparation of plants and *Agrobacterium* strains hosting the viral vectors.

***Alternatives***: The protocol uses viral vectors based on viruses naturally infecting species of the Solanaceae family, including *N. benthamiana*. Alternative plant species may be used depending on the known host range of the viral vectors and availability of transgenic lines expressing a Cas nuclease.

## Key resources table

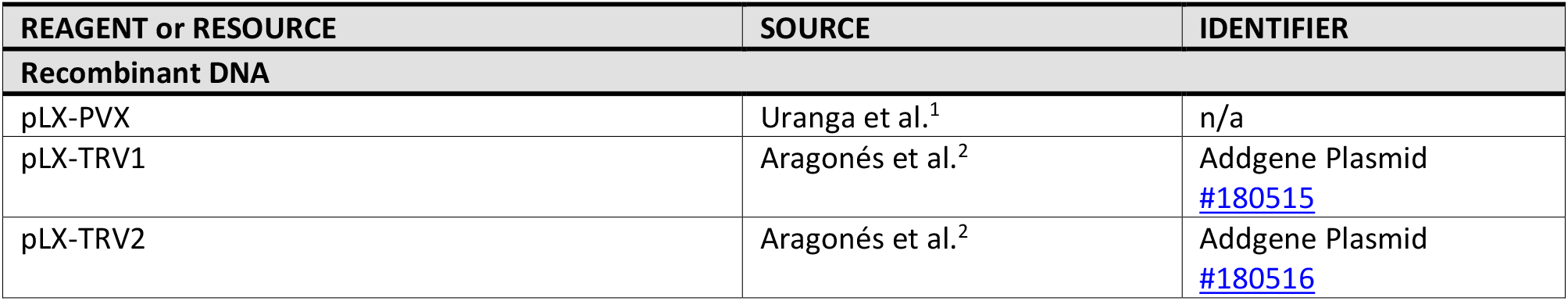
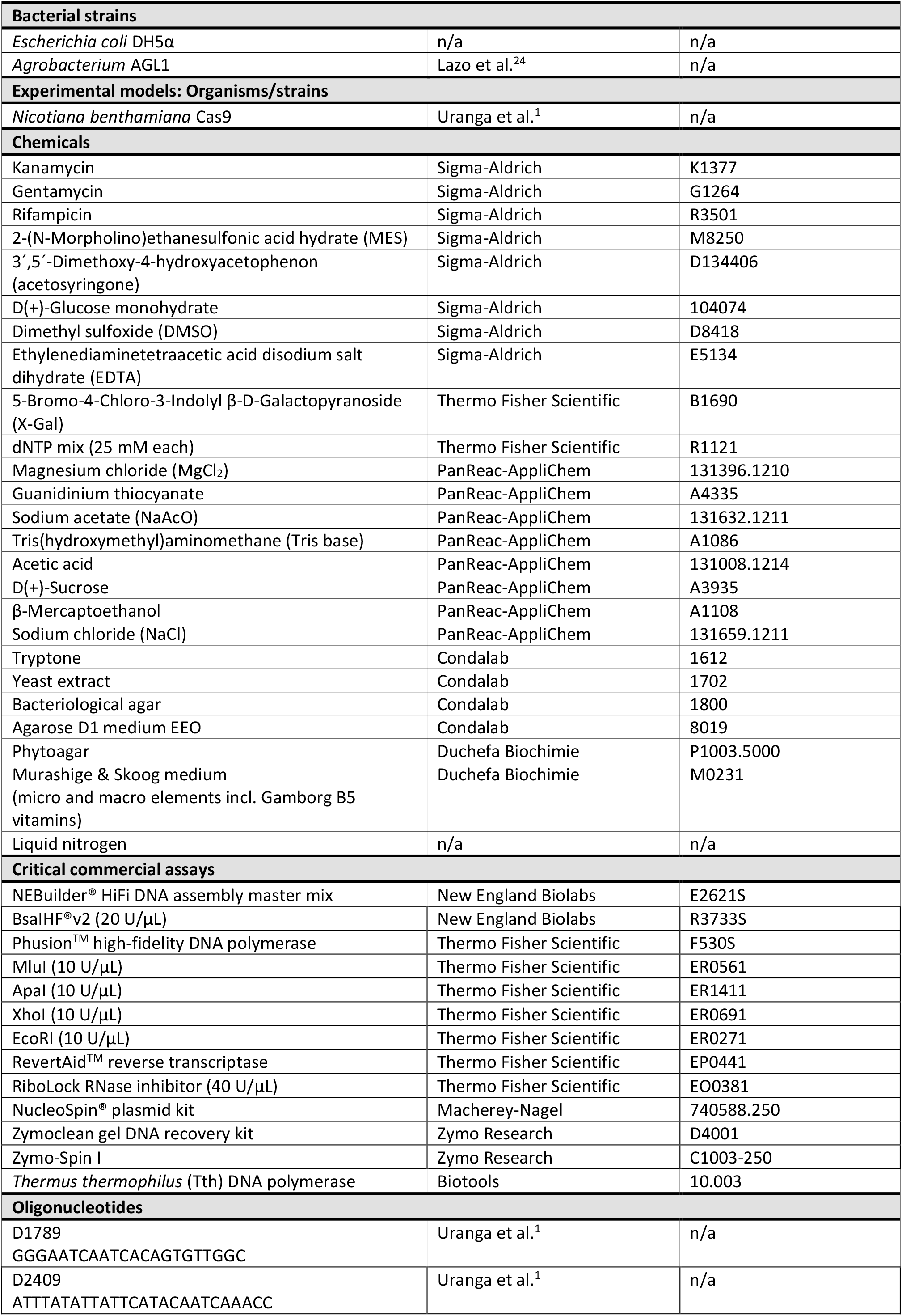
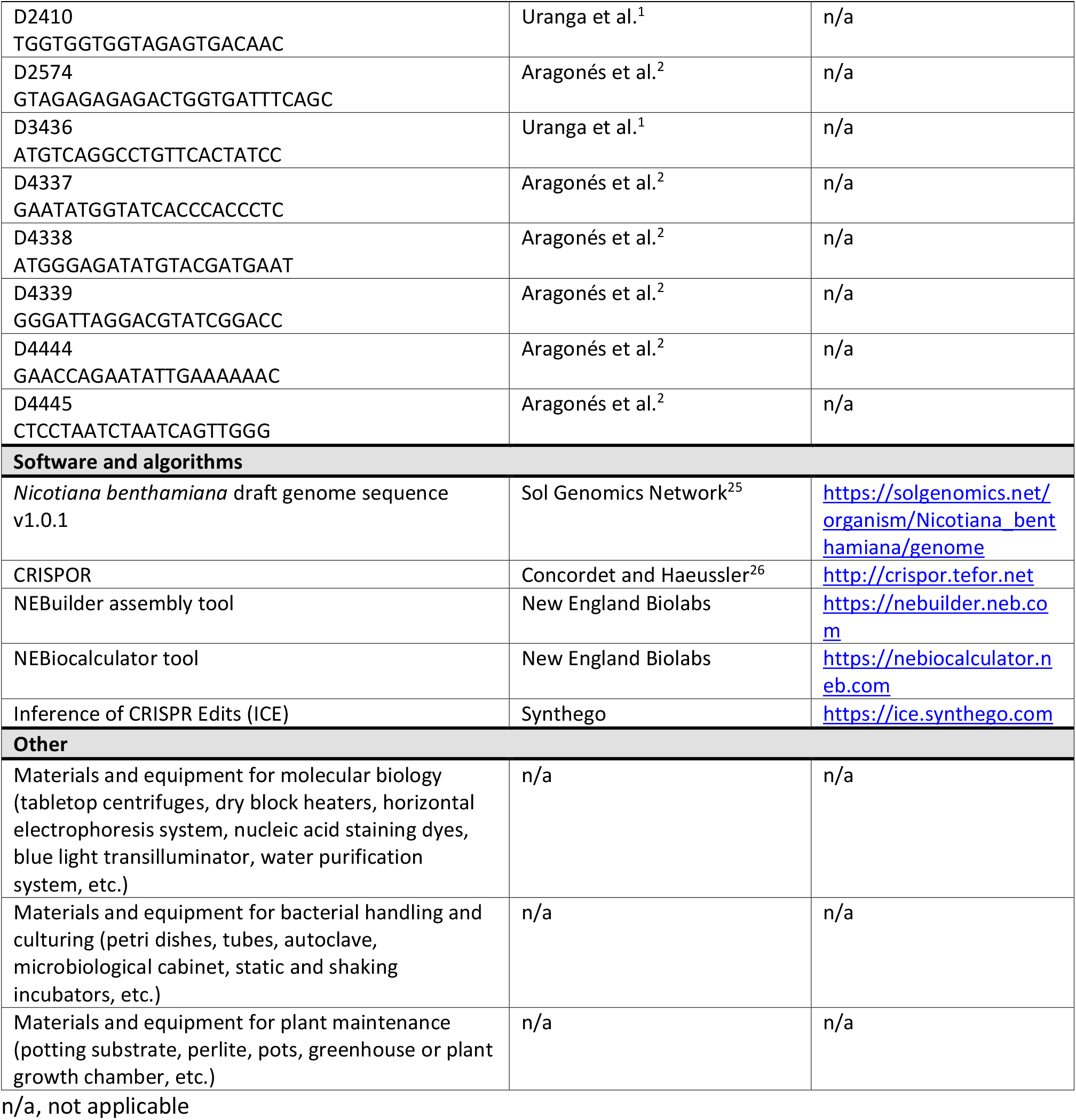

## Materials

***Note:*** Unless otherwise indicated, standard molecular cloning reagents and methods^27^ are used, and materials, solutions, media and buffers are prepared and stored at room temperature (20-25°C). Ultrapure water is obtained using a Milli-Q® system (Millipore), and autoclave sterilization is done at 121°C (20 min).

### Preparation of stock solutions and media

#### Timing: 1-2 days

- Kanamycin (50 mg/mL): Dissolve 0.5 g kanamycin powder in 10 mL ultrapure water, sterilize through a 0.2 μm filter, and store 1-mL aliquots at -20°C, for up to 1 year.
- Gentamycin (20 mg/mL): Dissolve 0.2 g gentamycin powder in 10 mL ultrapure water, sterilize through a 0.2 μm filter, and store 1-mL aliquots at -20°C, for up to 1 year.
- Rifampicin (50 mg/mL): Dissolve 0.5 g rifampicin powder in 10 mL DMSO, and store 1-mL aliquots at -20°C in the dark, for up to 6 months.
- Glucose (1 M): Dissolve 9.91 g of D(+)-glucose monohydrate in ultrapure water to a final volume of 50 mL, sterilize through a 0.2-μm filter, and store indefinitely.
- X-Gal (40 mg/mL): Dissolve 0.4 g X-Gal powder in 10 mL N, N-dimethylformamide. Store 1-mL aliquots at -20°C in the dark, for up to 6 months.
- NaAcO 3M: Dissolve 204.12 g NaAcO in 0.4 L ultrapure water, adjust pH to 5.5 with glacial acetic acid, take the volume up to 0.5 L with ultrapure water, and autoclave. Store for up to 1 year.
- MES pH 5.5 (1 M): Dissolve 9.762 g MES powder in 40 mL ultrapure water, adjust pH to 5.5 with 1 N KOH, take the volume up to 50 mL with ultrapure water, and sterilize through a 0.2 μm filter. Store in the dark, for up to 6 months.
- MgCl_2_ (2 M): Dissolve 9.52 g MgCl_2_ in ultrapure water to a final volume of 50 mL, and sterilize through a 0.2 μm filter. Store for up to 1 year.
- Acetosyringone (0.1 M): Dissolve 98 mg acetosyringone powder in 5 mL DMSO, and store in single-use 150-μL aliquots at -20°C in the dark, for up to 6 months.
- EDTA (0.5 M): Mix 93.05 g EDTA and 0.4 L distilled water on a magnetic stirrer, adjust pH to 8.0 with ∼10 g NaOH pellets (EDTA will not go into solution until the solution reach pH ∼8.0), take the volume up to 0.5 L with distilled water and autoclave. Store for up to 1 year.
- Oligonucleotides: Custom DNA oligonucleotides are purchased desalted after synthesis as 100 μM stock solutions. Stocks are diluted in autoclaved ultrapure water to have 10 μM aliquots for PCR, and 2 μM aliquots for cDNA synthesis.
- Media:

#### Lysogenic broth (LB)

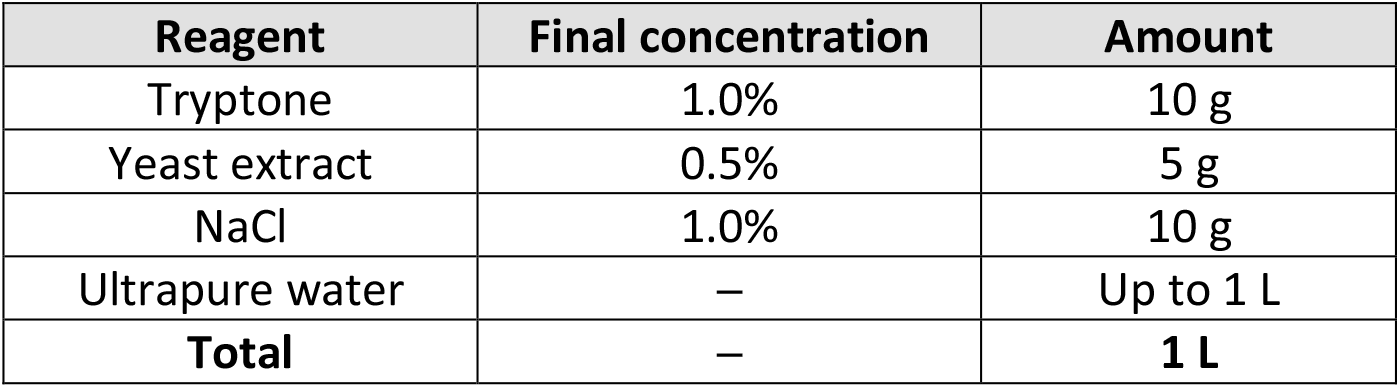
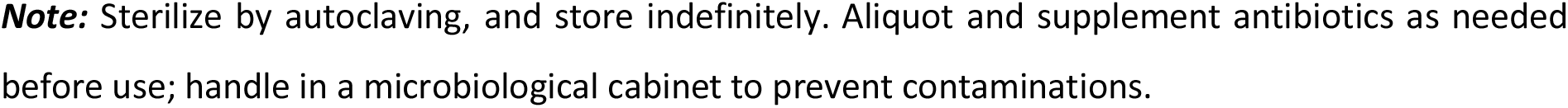

| Reagent | Final concentration | Amount |
| --- | --- | --- |
| Tryptone | 1.0% | 10 g |
| Yeast extract | 0.5% | 5 g |
| NaCl | 1.0% | 10 g |
| Ultrapure water | — | Up to 1 L |
| **Total** | — | **1 L** |

**Note:** Sterilize by autoclaving, and store indefinitely. Aliquot and supplement antibiotics as needed before use; handle in a microbiological cabinet to prevent contaminations.

#### LB agar

| Reagent | Final concentration | Amount |
| --- | --- | --- |
| Tryptone | 1.0% | 10 g |
| Yeast extract | 0.5% | 5 g |
| NaCl | 1.0% | 10 g |
| Bacteriological agar | 1.5% | 15 g |
| Ultrapure water | – | Up to 1 L |
| **Total** | – | **1 L** |

**Note:** Sterilize by autoclaving, cool the medium down to ~50–60°C and, in a microbiological cabinet, supplement antibiotics and mix vigorously. Aliquot into petri dishes, and once solidified store at 4°C in the dark.
**Note:** X-Gal solution can be spread onto LB agar plates before use.

#### Super optimal broth (SOB)

| Reagent | Final concentration | Amount |
| --- | --- | --- |
| Tryptone | 2.0% | 20 g |
| Yeast extract | 0.5% | 5 g |
| NaCl | 10.0 mM | 0.584 g |
| KCl | 2.5 mM | 0.186 g |
| MgSO<sub>4</sub> | 20.0 mM | 2.4 g |
| Ultrapure water | – | Up to 1 L |
| **Total** | – | **1 L** |

**Note:** Sterilize by autoclaving, and store indefinitely.
**Note:** Handle in a microbiological cabinet to prevent contaminations.

#### SOC

| Reagent | Final concentration | Amount |
| --- | --- | --- |
| SOB | – | 9.8 mL |
| Glucose (1 M) | 20 mM | 0.2 mL |
| **Total** | – | **10 mL** |

**Note:** Sterilize through a 0.2-µm filter and store indefinitely.
**Note:** Handle in a microbiological cabinet to prevent contaminations.

#### ½ Murashige & Skoog (MS)

| Reagent | Final concentration | Amount |
| --- | --- | --- |
| Murashige & Skoog medium | 0.5× | 2.15 g |
| (micro and macro elements incl. Gamborg B5 vitamins) |  |  |
| 2-(N-Morpholino)ethanesulfonic acid hydrate (MES) | 0.5 g/L | 0.5 g |
| Sucrose | 3% | 30 g |
| Phytoagar | 0.8% | 8 g |
| Ultrapure water | – | Up to 1 L |
| **Total** | – | **1 L** |

**Note:** Adjust to pH 5.7, sterilize by autoclaving, aliquot into petri dishes, and once solidified store at 4°C in the dark, for up to 6 months.

### Preparation of buffers

#### Timing: 1-2 days

##### Tris-acetate-EDTA (TAE) buffer (50×)

| Reagent | Final concentration | Amount |
| --- | --- | --- |
| Tris-base | 2 M | 242 g |
| EDTA pH 8 (0.5 M) | 50 mM | 100 mL |
| Acetic acid (100%) | 1 M | 57.1 mL |
| Ultrapure water | – | Up to 1 L |
| **Total** | – | **1 L** |

**Note:** Store indefinitely.

##### Infiltration buffer

| Reagent | Final concentration | Amount |
| --- | --- | --- |
| MES pH 5.5 (1 M) | 10 mM | 2.5 mL |
| MgCl<sub>2</sub> (2 M) | 10 mM | 1.25 mL |
| Acetosyringone (100 mM) | 150 µM | 0.375 mL |
| Ultrapure water | – | Up to 250 mL |
| **Total** | – | **250 mL** |

**Note:** Freshly prepared before use.

##### RNA/DNA extraction buffer

| Reagent | Final concentration | Amount |
| --- | --- | --- |
| Guanidinium thiocyanate | 4 M | 47.3 g |
| NaAcO pH 5.5 (3 M) | 0.1 M | 3.33 mL |
| EDTA pH 8.0 (0.5 M) | 10 mM | 2 mL |
| β-Mercaptoethanol | 0.1 M | 0.7 mL |
| Ultrapure water | – | Up to 100 mL |
| **Total** | – | **100 mL** |

**Note:** Freshly add β-Mercaptoethanol before use.

##### Washing buffer

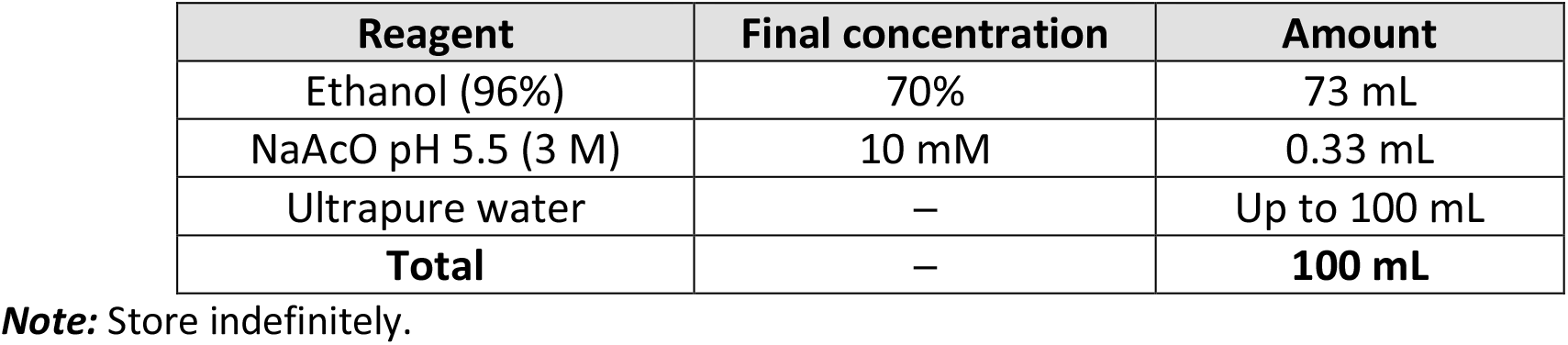

| Reagent | Final concentration | Amount |
| --- | --- | --- |
| Ethanol (96%) | 70% | 73 mL |
| NaAcO pH 5.5 (3 M) | 10 mM | 0.33 mL |
| Ultrapure water | – | Up to 100 mL |
| **Total** | – | **100 mL** |

**Note:** Store indefinitely.

##### Elution buffer

| Reagent | Final concentration | Amount |
| --- | --- | --- |
| Tris-HCl pH 8.5 (0.1 M) | 20 mM | 2 mL |
| Ultrapure water | – | Up to 10 mL |
| **Total** | – | **10 mL** |

**Note:** Sterilized by autoclaving and store indefinitely.

## Step-by-step method details

***Note:*** Unless otherwise indicated, standard molecular cloning reagents and methods^27^ are used, and protocol steps are done at room temperature (20-25°C).

### Homology-based assembly of recombinant viral vectors

#### Timing: 3-5 days

This section described the use of homology-based assembly to generate full-length clones of potato virus X (PVX; genus *Potexvirus*) and tobacco rattle virus (TRV; genus *Tobravirus*) expressing target-specific sgRNAs (Figure 2). Plasmids for recombinant viral vectors are built by standard molecular biology techniques, as described in Uranga et al.^1^ and Aragonés et al.^2^.

***Note:*** Simultaneous disruption of *Phytoene desaturase 3* (*PDS*) homeologs present in the genome of the allotetraploid *N. benthamiana* results in cells with a photobleaching phenotype typical of carotenoid biosynthetic defects^11,17,28,29^. *PDS* is used in the described procedures as the target gene to facilitate visual inspection of CRISPR-Cas9-mediated genome editing in somatic cells and progeny.

1. sgRNA design:
  a. Download the cDNA sequence of the target *N. benthamiana* gene from Sol Genomics Network (https://solgenomics.net/organism/Nicotiana_benthamiana/genome/). ***Note:*** Download Niben101Scf01283g02002.1 and Niben101Scf14708g00023.1 sequences for targeting of *PDS* homeologs
  b. Paste the cDNA sequence into CRISPOR bioinformatics software (http://crispor.tefor.net/) and set the following parameters:
    i. *N. benthamiana* – SolGenomics.net V1.01 as the genome
    ii. 20 bp-NGG – SpCas9, SpCas9-HF1, eSpCas9 1.1 as the Protospacer Adjacent Motif (PAM)
  c. Select one or two protospacers that target the coding region of a gene and show a predicted efficiency of ≥ 45 (i.e. Doench ‘16 and Mor.-Mateos scores). ***CRITICAL:*** To increase the chance to recover knock-out mutants, avoid protospacers that target splicing junctions and non-coding regions (i.e. introns, 5’ and 3’ UTRs). ***CRITICAL:*** *N. benthamiana* genome is allotetraploid and loss-of-function mutations of two homeologs may be required to obtain a mutant phenotype. ***Note:*** Select the protospacer TTGGTAGTAGCGACTCCATG conserved in Niben101Scf01283g02002.1 and Niben101Scf14708g00023.1 for simultaneous targeting and disruption of two *PDS* homeologs (Figure 3A).

**Figure 3.**
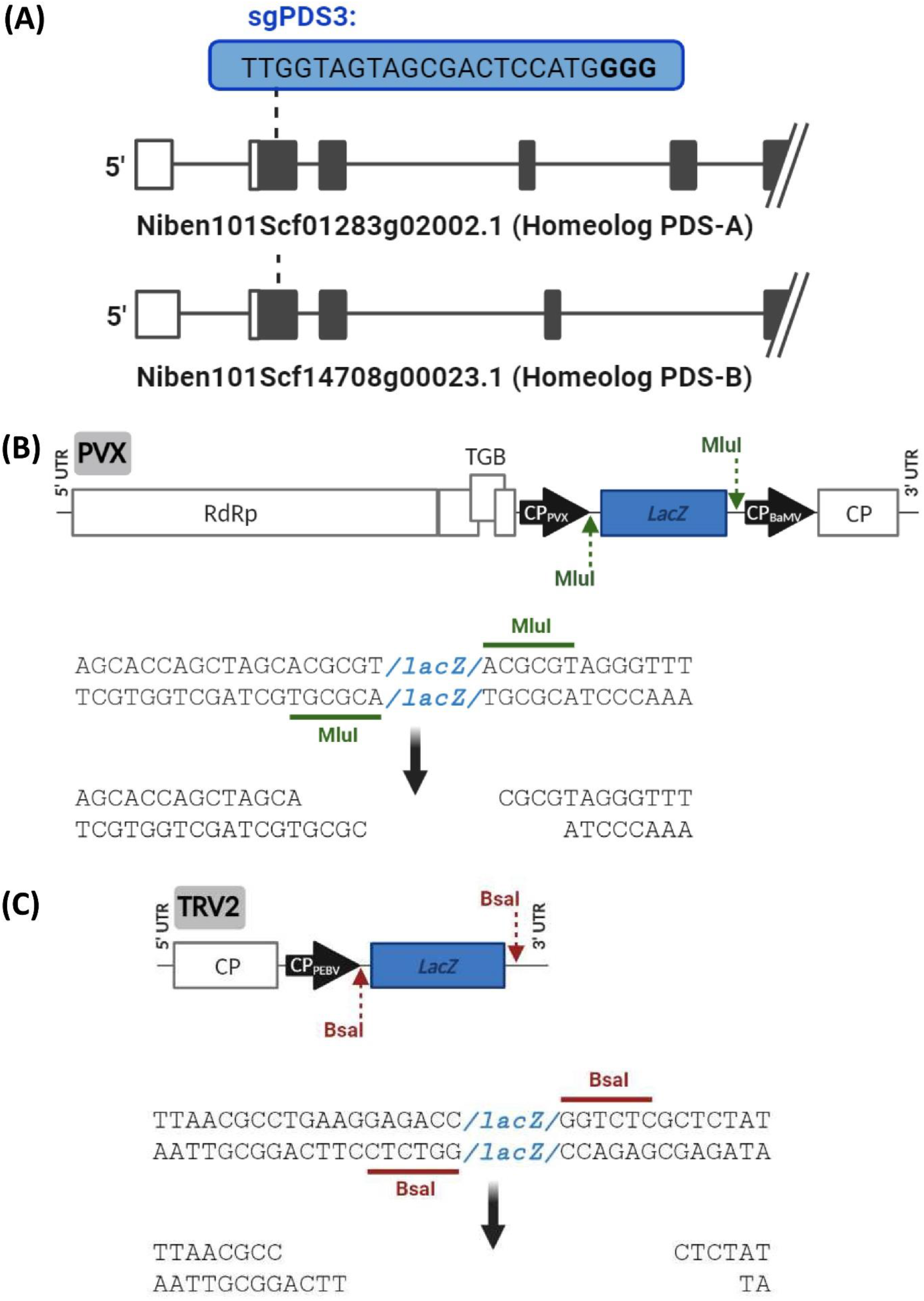
Features of pLX-PVX and pLX-TRV2 cloning cassettes for sgRNA delivery. **(A)** The protospacer of sgPDS3-tFT is designed to target a 20-nt conserved region immediately downstream the starting codon of two *N. benthamiana PDS* homeologs (Niben101Scf01283g02002.1 and Niben101Scf14708g00023.1). **(B)** Schematic representation of pLX-PVX cloning cassette is shown along with the MluI recognition sites (green underline) and MluI-generated overhangs (bottom). The *LacZ* reporter (blue box) allows white-blue screen of recombinant vectors. Plant expression of the insert is driven by the endogenous coat protein (CP) promoter. **(C)** Schematic representation of pLX-TRV2 cloning cassette is shown along with the BsaI recognition sites (red underline) and BsaI-generated overhangs (bottom). Plant expression of the insert is driven by the pea early browning virus (PEBV) CP promoter. Other details are as in panel B.
  d. Downstream the selected protospacer(s) append the *S. pyogenes* Cas9 binding scaffold sequence: GTTTTAGAGCTAGAAATAGCAAGTTAAAATAAGGCTAGTCCGTTATCAACTTGAAAAAGTGGCACCGAGTCGGTGC
  e. Downstream the obtained sgRNA sequence append an RNA mobility element consisting of the first 102 bp of the *Arabidopsis thaliana Flowering locus T* (tFT; TAIR ID: AT1G65480) coding region: ATGTCTATAAATATAAGAGACCCTCTTATAGTAAGCAGAGTTGTTGGAGACGTTCTTGATCCGTTTAATAGATCAATCACTCTAAAGGTTACTTATGGCCAA ***CRITICAL:*** The truncated *A. thaliana* FT (tFT) sequence was shown to promote the movement of viral RNAs into the meristem^30^, and the sgRNA-tFT fusion greatly increased the frequencies of somatic and germline mutations in *N. benthamiana*^11^.
  f. Flank the obtained sgRNA-tFT sequence with cloning adapters suitable for homology-based assembly. ***Note:*** To assemble sgRNA-tFT into pLX-PVX, append the sequence GAGGTCAGCACCAGCTAGCA to the 5’ terminus, and TAGGGTTTGTTAAGTTTCCC to the 3’ terminus of the sequence from the previous step. ***Note:*** To assemble sgRNA-tFT into pLX-TRV2, append the sequence CACTTACCCGAGTTAACGCC to the 5’ terminus, and ATGTCCCGAAGACATTAAAC to the 3’ terminus of the sequence from the previous step. ***Note:*** Sequences of the fragments sgPDS3-PVX and sgPDS3-TRV2 for homology-based cloning of a sgRNA-tFT that targets *PDS* into pLX-PVX or pLX-TRV2, respectively, are as follows:

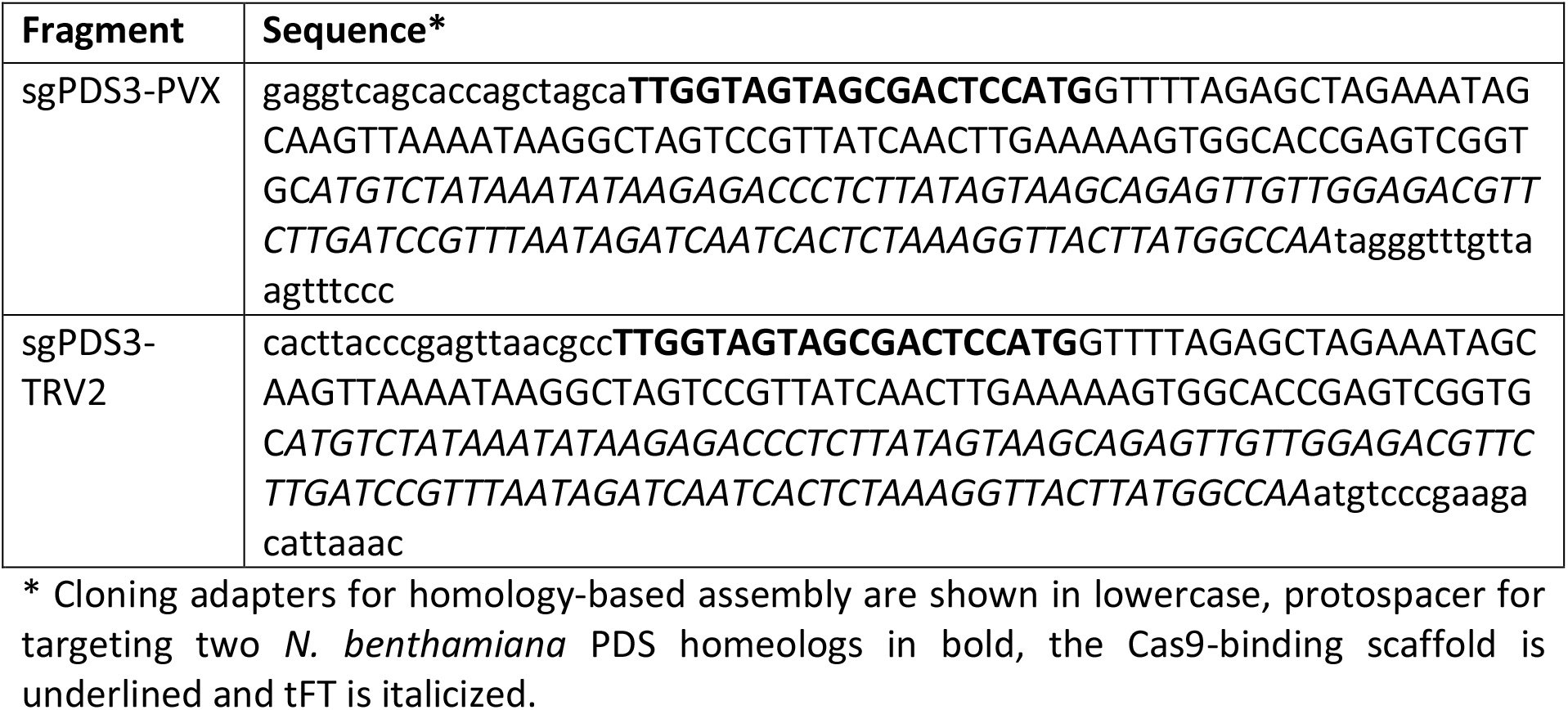
2. *In silico* assembly of recombinant virus clones. ***Note***: NEBuilder assembly tool (https://nebuilder.neb.com/) and Benchling bioinformatics tool (https://benchling.com/) can be used for the assembly simulation.
  a. For sgRNA-tFT assembly into pLX-PVX (Figure 3B)
    i. Retrieve the pLX-PVX sequence (see Supporting Information)
    ii. Perform an *in silico* restriction digest with MluI to linearize pLX-PVX ***Note:*** Digestion results in a 9.9-kb linearized fragment without the *lacZ* region.
    iii. Simulate the homology-based assembly reaction including the linearized vector and the linear sgRNA-tFT fragment designed in step 1f. ***Note:*** sgPDS3-PVX is used for *PDS* targeting
  b. For sgRNA-tFT assembly into pLX-TRV2 (Figure 3C)
    i. Retrieve the pLX-TRV2 sequence (GenBank: OM372496)
    ii. Perform an *in silico* restriction digest with BsaI-HF to linearize pLX-TRV2 ***Note:*** Digestion results in a 5.8-kb linearized fragment without the lacZ region.
    iii. Simulate the homology-based assembly reaction including the linearized vector and the linear sgRNA-tFT fragment designed in step 1f. ***Note:*** sgPDS3-TRV2 is used for *PDS* targeting
3. Chemically synthesize a linear double-stranded DNA fragment corresponding to the sgRNA-tFT sequence designed in step 1.f and resuspend it in ultrapure water to ∼25 ng/μL. ***Note:*** Chemically synthesized fragments are purchased as gBlocks™ Gene Fragment (Integrated DNA Technologies). ***Alternatives:*** Custom DNA fragments from alternative commercial providers may be used (e.g. GeneArt™ Strings DNA Fragments, Thermo Fisher Scientific; Twist Gene Fragments without adapters, Twist Bioscience; etc.). ***Pause point:*** Resuspended DNA fragments can be frozen and stored at -20°C several days.
4. pLX vector backbone preparation (Figure 3B, C):
  a. Linearize the viral vectors by restriction digestion and incubate at 37°C (1 h):

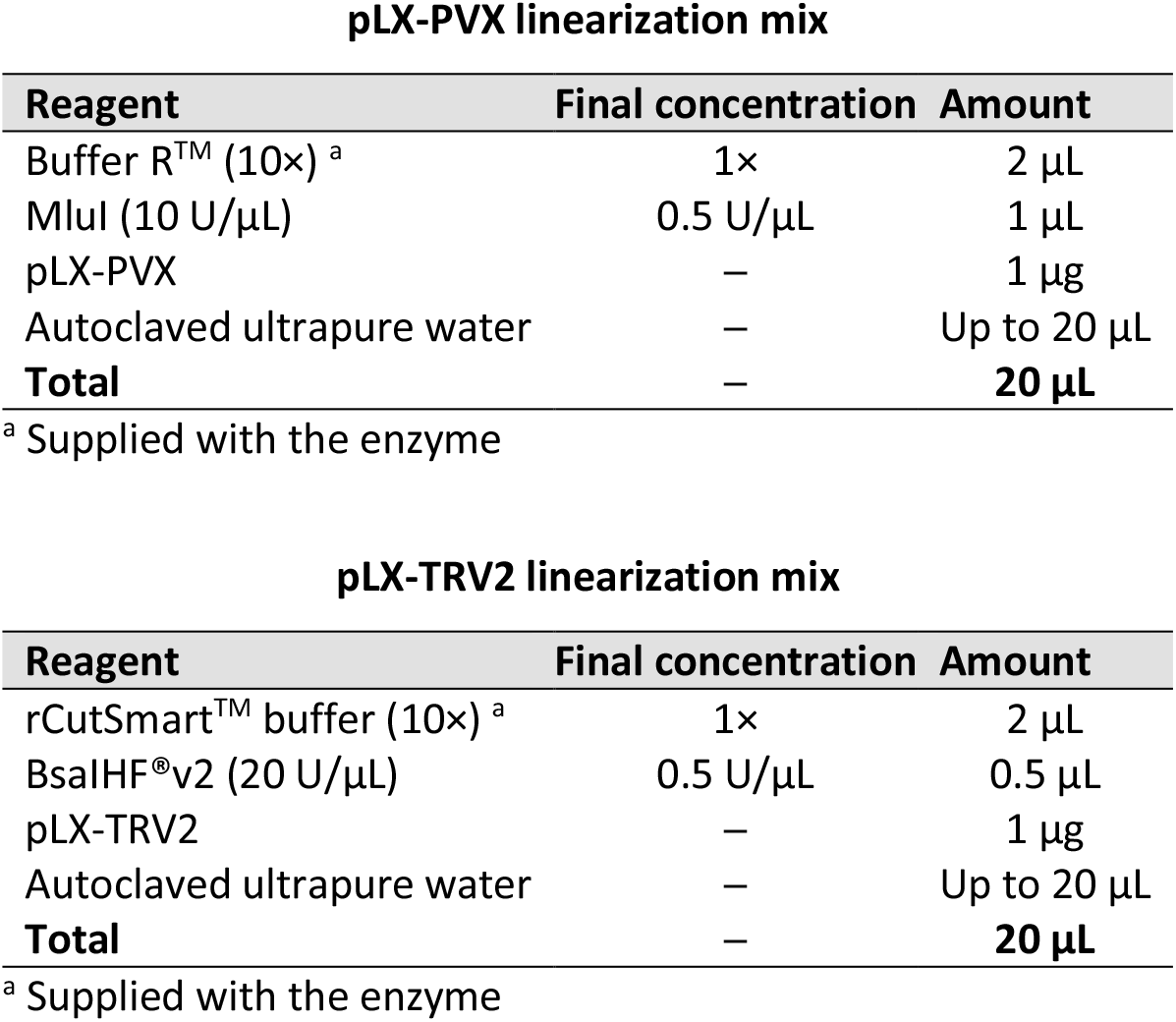 ***Pause point:*** Reaction can be frozen and stored at -20°C several days.
  b. Prepare agarose gel in 1× TAE buffer and run electrophoresis of digestion reactions. ***Note:*** Agarose D1 medium EEO is used for gel preparation; 0.8% agarose gels provide good electrophoresis resolution of the large (5-10 kb) DNA fragments.
  c. Cut bands on a blue light transilluminator using single-use disposable scalpels. ***Note:*** If a blue light transilluminator is not available a UV light transilluminator can be used but ensure to avoid prolonged exposure to UV light as it damages DNA samples and dramatically reduces assembly efficiency. If UV light transilluminator is used, wear suitable protection personal protective equipment to avoid UV-associated hazards. ***Pause point:*** Harvested gel bands can be frozen and stored at -20°C several days.
  d. Purify DNA from gel bands using the Zymoclean gel DNA recovery kit (Zymo Research) as per manufacturer’s instructions. ***Note:*** Other commercial gel purification kits can also be used. ***Pause point:*** Gel-purified DNA can be frozen and stored at -20°C several days.
5. Resolve by agarose gel electrophoresis 1 μL of the synthetic DNA fragments (step 3) and the gel-purified linearized vector (step 4) to check integrity, and estimate DNA amount and quality using a spectrophotometer (e.g. NanoDrop™, Thermo Fisher Scientific). ***Note:*** Use for downstream applications DNA samples that show a sharp single band, and with a ratio of absorbance at 260 nm and 280 nm (260/280) of ∼1.8, and 260/230 ratio in the range of 2.0-2.2.
6. *In vitro* homology-based assembly:
  a. Mix the gel-purified linearized vector (step 4) and a synthetic DNA fragment (step 3) in a molar ratio of 1:2 and a total volume of ∼2μL. ***Note:*** Use a total DNA amount of 0.05-2 pmol. ***Note:*** The NEBiocalculator® Tool (https://nebiocalculator.neb.com/) can be used for molarity calculations.
  b. Add 1 volume (∼2 μL) of NEBuilder® HiFi DNA assembly master mix (New England Biolabs). ***Alternative:*** Gibson assembly^31^ can also be used for homology-based cloning; the required enzymatic mix can be in-house prepared as detailed^32^, or purchased from commercial providers (e.g. Gibson Assembly® master mix, New England Biolabs, cat. number E2611S). If using Gibson assembly, no protocol changes are required.
  c. Incubate the mix at 50°C (1 h). Once the reaction ends, place tubes on ice.
7. Transformation of *E. coli* cells:
  a. Thaw a 40-μL aliquot of electrocompetent *E. coli* DH5α cells on ice. ***Alternative:*** Other *E. coli* strains suitable for high-efficient transformation may be used.
  b. Gently mix by pipetting a cell aliquot with 1 μL of the assembly reaction from step 6.b, and transfer the mixture to a pre-chilled electroporation cuvette.
  c. Electroporate cells (1500 V), mix them with 1 mL of room-temperature SOC (see Materials) and transfer to a 1.5-mL tube. ***Note:*** Once DNA is added to cells electroporation can be carried out immediately, without any incubation. ***Note:*** If your sample arcs when electroporating, dialyze or purify the assembly reaction to remove salt excess. ***Alternative:*** Heat shock of chemically-competent *E. coli* cells for high-efficiency transformation^23^ may be used.
  d. Incubate the cells at 37°C, shaking vigorously at 200-250 rpm (1 h). ***Note:*** Warm selection plates at 37°C while cells are recovering.
  e. Pellet bacteria by centrifuging at 15 000 × g (1 min), resuspend bacteria in 100 μL of SOC and plate the suspension onto LB agar plates supplemented with 50 mg/L kanamycin and 40 μL X-Gal (40 mg/mL). Incubate at 37°C (overnight). ***Note:*** Plate the X-Gal solution on agar plates before use for white-blue screen of recombinant clones. ***Note:*** Bacterial cells harboring binary vectors with full-length virus clones generally display low growth rates; plate incubation up to 24 h may be necessary. ***Pause point:*** After colony appearance, plates can be stored 1-3 days at 4°C.
8. Confirm the presence of recombinant binary vectors in *E. coli* transformants:
  a. Pick a white colony obtained by X-Gal selection and inoculate 5-10 mL liquid LB with 50 mg/L kanamycin in 50 mL tubes. Incubate the tubes at 37°C, shaking vigorously at 200-250 rpm (overnight). ***Note:*** Bacterial cells harboring binary vectors with full-length virus clones generally display low growth rates; extend culturing time up to 24 h if necessary. ***Note:*** It is recommended to purify plasmid DNA from 2–4 colonies per construct.
  b. Harvest bacteria from the culture by centrifugation at 15 000 × g (2 min); discard the medium, centrifuge again (30 s) and remove medium residues by pipetting. ***Pause point:*** Harvested bacterial pellets can be frozen and stored at -80°C several days before proceeding with plasmid DNA purification.
  c. Purify DNA plasmids from bacterial pellets using NucleoSpin® plasmid kit (Macherey Nagel) as per manufacturer’s instructions, except that:
    i. Use double volumes of resuspension, lysis and neutralization solutions supplied within the kit to improve clearing of bacterial lysates and increase the amount of purified plasmid. ***Note:*** After addition of the lysis solution incubate until the cell suspension clears (5 min).
    ii. Transfer ∼800 μL of cleared lysate to the column, centrifuge, discard the flowthrough and load again the same column with the remainder of the lysate, ***Note:*** The volume of the cleared bacterial lysate will exceed the spin column capacity. ***Note:*** Other commercial plasmid DNA purification kits can also be used. ***Pause point:*** Purified DNA plasmids can be frozen and stored at -20°C indefinitely.
9. Identify recombinant viral vectors by restriction enzyme digestion of purified DNA plasmids (37°C, 1 h), and agarose gel electrophoresis.

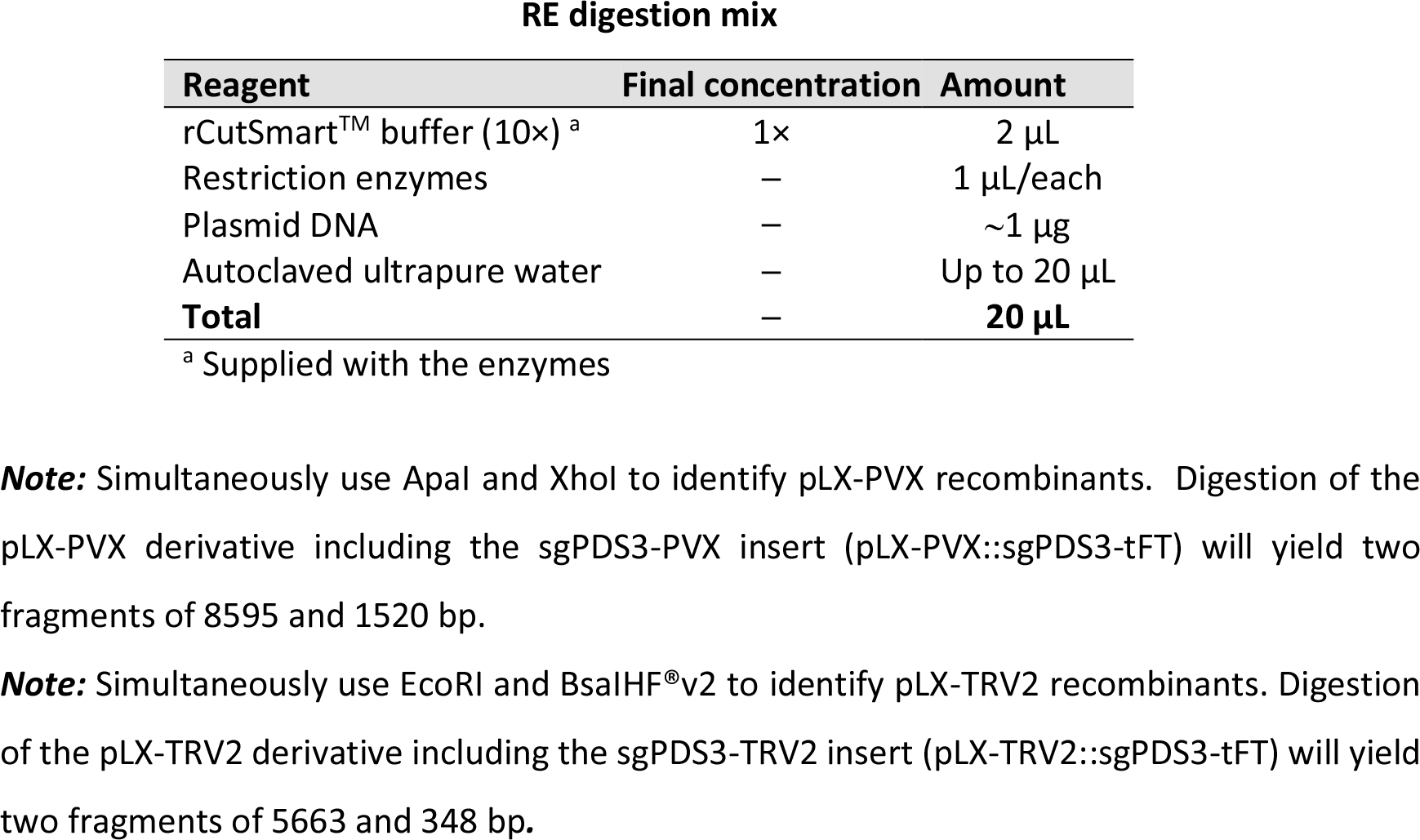 ***Pause point:*** DNA plasmids with the correct digestion profiles can be frozen and stored at - 20°C indefinitely.
10. Verify the insert sequence of the identified recombinant viral vectors by Sanger sequencing. ***Note:*** Use primer D1789 to sequence pLX-PVX inserts. ***Note:*** Use primer D2574 to sequence pLX-TRV2 inserts. ***Note:*** It is recommended to sequence 2 clones per construct.

### *Agrobacterium*-mediated inoculation of plants

#### Timing: 4-7 days

In this section, binary vectors with the correct sequence confirmed by sequencing are transformed into *Agrobacterium* AGL1 cells. Selected cells harboring the binary vectors are then cultured, prepared for inoculation and used to inoculate *N. benthamiana* Cas9 plants.

***CRITICAL:*** Ensure to coordinate preparation of *N. benthamiana* plants and *Agrobacterium* strains hosting the viral vectors.

***CRITICAL:*** JoinTRV relies on two T-DNA vectors with compatible origins for simultaneous agroinoculation of TRV genomic components^2^. To obtain a single *Agrobacterium* strain co-hosting the JoinTRV vectors, *Agrobacterium* AGL1 is first transformed with pLX-TRV1 and selected on plates supplemented with 20 mg/L gentamycin and 50 mg/L rifampicin; competent cells of the recovered AGL1 pLX-TRV1 strain are then prepared and used in step 11.

***Alternative:*** JoinTRV vectors have autonomous replication and can be used in a one-vector/one-*Agrobacterium* approach. In this case: (i) transform pLX-TRV1 into *Agrobacterium* cells and selected on plates supplemented with 20 mg/L gentamycin and 50 mg/L rifampicin (step 11); (ii) transform pLX-TRV2 into *Agrobacterium* cells and selected on plates supplemented with 50 mg/L kanamycin and 50 mg/L rifampicin (step 11); (iii) individually culture the two obtained *Agrobacterium* strains as described in steps 12-14 and pool them before to proceed with the preparation of bacterial suspensions for inoculation (step 15).

11. Transformation of *Agrobacterium* cells:
  a. Thaw aliquots of electrocompetent *Agrobacterium* AGL1 cells on ice. ***Alternatives:*** Viral vectors of this protocol are based on binary T-DNA vectors of the pLX series, which have been tested in a variety of *Agrobacterium* strains^21^. Strain alternatives include C58C1-313, EHA105, CryX, NMX021 (ref. ^20,33–35^), as well as any *Agrobacterium* strain capable of plant cell transformation and sensitive to kanamycin and gentamicin, the antibiotic used for pLX-PVX, pLX-TRV1 and pLX-TRV2 selection. ***CRITICAL:*** GV3101::pMP90 is a common *Agrobacterium* strain resistant to gentamicin^35^ not compatible with the JoinTRV system.
  b. Gently mix by pipetting a cell aliquot with ∼0.5 μg of plasmid DNA, and transfer the mixture to a pre-chilled electroporation cuvette.
    i. Use standard AGL1 cells for pLX-PVX transformation.
    ii. Use cells of the AGL1 pLX-TRV1 strain for pLX-TRV2 transformation. ***Note:*** It is recommended to independently transform *Agrobacterium* cells with plasmid DNA obtained from two *E. coli* colonies per construct. ***Note:*** If your sample arcs when electroporating, dilute or purify plasmid DNA to remove salt excess.
  c. Electroporate cells (1500 V), mix them with 1 mL of room-temperature SOC and transfer to a 1.5-mL tube. ***Note:*** Once DNA is added to cells electroporation can be carried out immediately, without any incubation. ***Alternative:*** Freeze-thaw transformation^23^ of *Agrobacterium* cells may be used.
  d. Incubate the cells at 28°C, shaking vigorously at 200-250 rpm (2-4 h). ***Note:*** Warm selection plates at 28°C while cells are recovering.
  e. Pellet bacteria by centrifuging at 15 000 × g (1 min), resuspend bacteria in 100 μL of SOC and plate the suspension onto LB agar plates supplemented with antibiotics.
    i. Use LB agar plates with 50 mg/L rifampicin and 50 mg/L kanamycin for pLX-PVX selection.
    ii. Use LB agar plates with 50 mg/L rifampicin, 20 mg/L gentamycin and 50 mg/L kanamycin for simultaneous selection of pLX-TRV1 and pLX-TRV2.
  f. Incubate plates at 28°C (48-72 h). ***Note:*** Bacterial cells harboring binary vectors with viral vectors may display low growth rates; extended plate incubation time may be necessary. ***Pause point:*** After colony appearance, plates can be stored 1-3 days at 4°C.
12. Culturing of *Agrobacterium* strains harboring the viral vectors
  a. Prepare 12 mL tubes with 3 mL liquid LB supplemented with antibiotics
    i. Use 50 mg/L kanamycin and 50 mg/L rifampicin for pLX-PVX selection
    ii. Use 20 mg/L gentamycin, 50 mg/L kanamycin and 50 mg/L rifampicin for JoinTRV selection (pLX-TRV1/pLX-TRV2).
  b. Inoculate tubes with individual colonies from step 11. ***CRITICAL:*** Bacteria harboring viral vectors may display low growth rates. If a mixture of small and large colonies appears during agar plate selection of transformed bacteria pick small colonies for subsequent analysis.
  c. Incubate tubes at 28°C, shaking at 200-250 rpm (24-48 h).
13. Use 0.7-mL culture aliquots from the previous step to prepare 20% glycerol bacterial stocks and store them at -80°C indefinitely.
14. Use 0.1-mL culture aliquots from step 12 to inoculate 10 mL liquid LB with (for pLX-PVX) or with 20 mg/L gentamycin and 50 mg/L kanamycin (for pLX-TRV1/pLX-TRV2) in 50 mL tubes. Incubate tubes at 28°C, shaking at 200-250 rpm (overnight).
15. Preparation of bacterial suspensions for inoculation:
  a. Measure in a spectrophotometer the optical density at 600 nm (OD_600_) of the bacterial suspensions until it reaches an OD_600_ ≈ 1.
  b. Pellet bacteria by centrifugation at 7200 × g (5 min), discard the medium, centrifuge again (30 s) and remove medium residues by pipetting.
  c. Resuspend the cell pellet to an OD_600_ of 0.5 in 4 mL Infiltration Buffer.
  d. Incubate at 28°C in the dark (2 h) for induction of bacterial genes required for T-DNA delivery to plant cells.
16. Plant inoculation:
  a. Water plants before starting with the inoculation to ensure they are well hydrated and induce stomata opening. ***Note:*** Organize and label plants before starting the inoculation. ***Note:*** Include mock-inoculated and wild-type plants as controls.
  b. Inoculate two young leaves per plant on the abaxial side using a 1-mL needleless syringe. Infiltrate ∼1-2 mL of bacterial suspension per leaf. ***CRITICAL:*** Make sure not to press the needleless syringe so hard as to pierce the leaf through.
  c. Transfer the plants to a growth chamber set at 25°C and 12 h-light/12 h-dark cycle. ***Note:*** To avoid cross-contaminations, ensure that plants infected with different constructs do not touch each other and they do not share the same watering tray. ***Note:*** Other growth chamber settings may work.

### Analysis of plant virus infectivity

#### Timing: 14-21 days

In this section, plants inoculated with full-length virus clones are monitored for the appearance and development of infection symptoms (Video 1). Biochemical assays are used to confirm the presence of virus within plant tissues and analyze CRISPR-Cas-mediated genome editing.

17. Inspect inoculated plants every 2-3 days and record appearance of infection symptoms. ***Note:*** *N. benthamiana* plants inoculated with pLX-PVX will display symptoms characterized by vein banding, ring spots and leaf atrophy starting after ∼1 week. Virus infection started by simultaneous inoculation of pLX-TRV1 and pLX-TRV2 can be very mild or symptomless. ***Note:*** Inoculation of Cas9-transgenic plants with viral vectors with sgRNA-tFT targeting the *PDS* homeologs will result in a leaf mosaic with green and photobleached sectors starting after ∼2 weeks (Video 1).
18. Analysis of virus infectivity ***CRITICAL:*** To prevent false positive results, avoid contaminations with plasmid DNA of the viral vectors during RNA extraction and the RT-PCR assays.
  a. Collect samples of the first symptomatic systemic leaf with a 1.2-cm cork borer (∼100 mg of tissue) at 14 days post-infiltration (dpi) and immediately freeze in liquid nitrogen. ***CRITICAL:*** To prevent false positives, only upper uninoculated plant samples should be analyzed and agrobacteria contamination avoided. ***CRITICAL:*** Collect samples from the third or younger leaves above the inoculated leaves. ***CRITICAL:*** Collect samples from mock-inoculated plants to be used as negative controls of virus detection assays. ***Pause point:*** After collection, samples can be stored at -80°C indefinitely.
  b. Total RNA extraction:
    i. Homogenize frozen leaf samples using a ball mill (Star-Beater, VWR) at 30 Hz (1 min).
    ii. Add 1 mL of RNA/DNA extraction buffer (see Materials), vortex (30 s) and centrifuge at 15,000 × g (5 min).
    iii. Transfer 0.6 mL of the supernatant to a 2-mL tube and add 0.39 mL (0.65 vol) of 96% ethanol, vortex (30 s), and centrifuge at 15,000 × g (1 min).
    iv. Transfer 0.7 mL of the supernatant to a silica column (Zymo-Spin I, Zymo Research), centrifuge at 15,000 × g (1 min) and wash twice with 0.5 mL of Washing buffer (see Materials).
    v. Centrifuge again (2 min) to remove buffer residues.
    vi. Transfer the silica column to a 1.5-mL tube, add 10 μL of 20 mM Tris-HCl pH 8.5 and incubate for 1 min, finally recover the eluted RNA sample by centrifugation at 15,000 × g (1 min).
    vii. Estimate RNA amount and quality using a spectrophotometer (e.g. NanoDrop™, Thermo Fisher Scientific). ***Note:*** Use for downstream applications RNA samples with a ratio of absorbance at 260 nm and 280 nm (260/280) of ∼2.0, and 260/230 ratio in the range of 2.0-2.2. ***Alternative:*** Other procedures and commercial kits for plant total RNA purification can also be used. ***Pause point:*** RNA samples can be stored for weeks at -80°C.
  c. Subject 1 μg of total RNA to cDNA synthesis using virus-specific primers and the RevertAid™ reverse transcriptase kit including RiboLock RNase inhibitor (Thermo Fisher Scientific) as per manufacturer’s instructions. ***Note:*** Use primer D2409 for PVX samples. ***Note:*** Use primer D4337 for TRV samples. ***Pause point:*** Reactions can be stored for several days at -20°C.
  d. Subject cDNA samples to PCR amplification using virus-specific primers.

Tth PCR master mix
| Reagent | Final concentration | Amount |
| --- | --- | --- |
| 10× Standard reaction buffer (with MgCl<sub>2</sub>)<sup>a</sup> | 1× | 2 µL |
| dNTPs (10 mM each) | 200 µM of each | 0.4 µL |
| Forward primer (10 µM) | 0.5 µM | 1 µL |
| Reverse primer (10 µM) | 0.5 µM | 1 µL |
| Tth DNA polymerase (1 U/µL) | 0.02 U/µL | 0.2 µL |
| cDNA sample | – | 1 µL |
| Ultrapure water | – | Up to 20 µL |
| **Total** | – | **20 µL** |
| <sup>a</sup> Supplied with the enzyme |  |  |

**Note:** Use the primer pair D3436/D2410 to amplify a 627-nt region of the PVX genome.
**Note:** Use the primer pair D4338/D4339 to amplify a 612-nt region of the TRV genome.

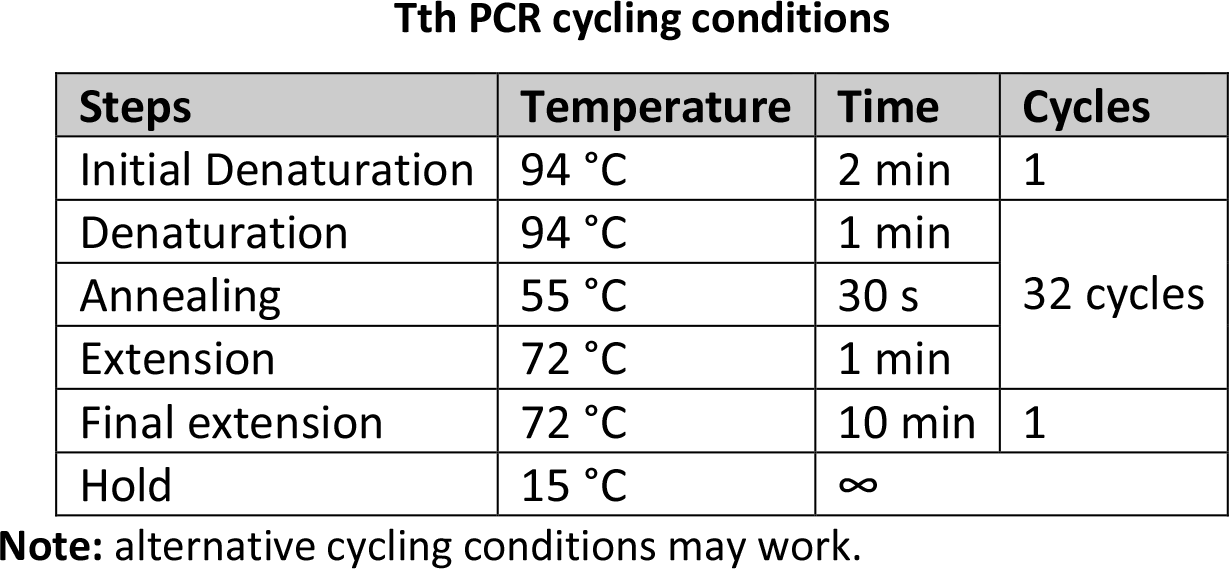 ***Pause point:*** Reactions can be stored for several days at -20°C.
  e. Analyze PCR products by electrophoresis in 1% agarose gels in 1× TAE buffer. ***Note:*** 627-bp and 612-bp amplicons are respectively detected in samples inoculated with the viral vectors based on PVX and TRV.

### Analysis genome editing of somatic cells

#### Timing: 2-4 weeks

In this section, biochemical assays are used to analyze CRISPR-Cas-mediated genome editing in inoculated plants.

19. Detection of Cas9-mediated genome editing in somatic cells of inoculated plants (Figure 4, Video 1):
  a. Collect samples of the first symptomatic systemic leaf with a 1.2-cm cork borer (∼100 mg of tissue) at 14 and 21 dpi and immediately freeze in liquid nitrogen. ***CRITICAL:*** Collect samples from mock-inoculated or wild-type plants to be used as negative controls of editing detection assays.

**Figure 4.**
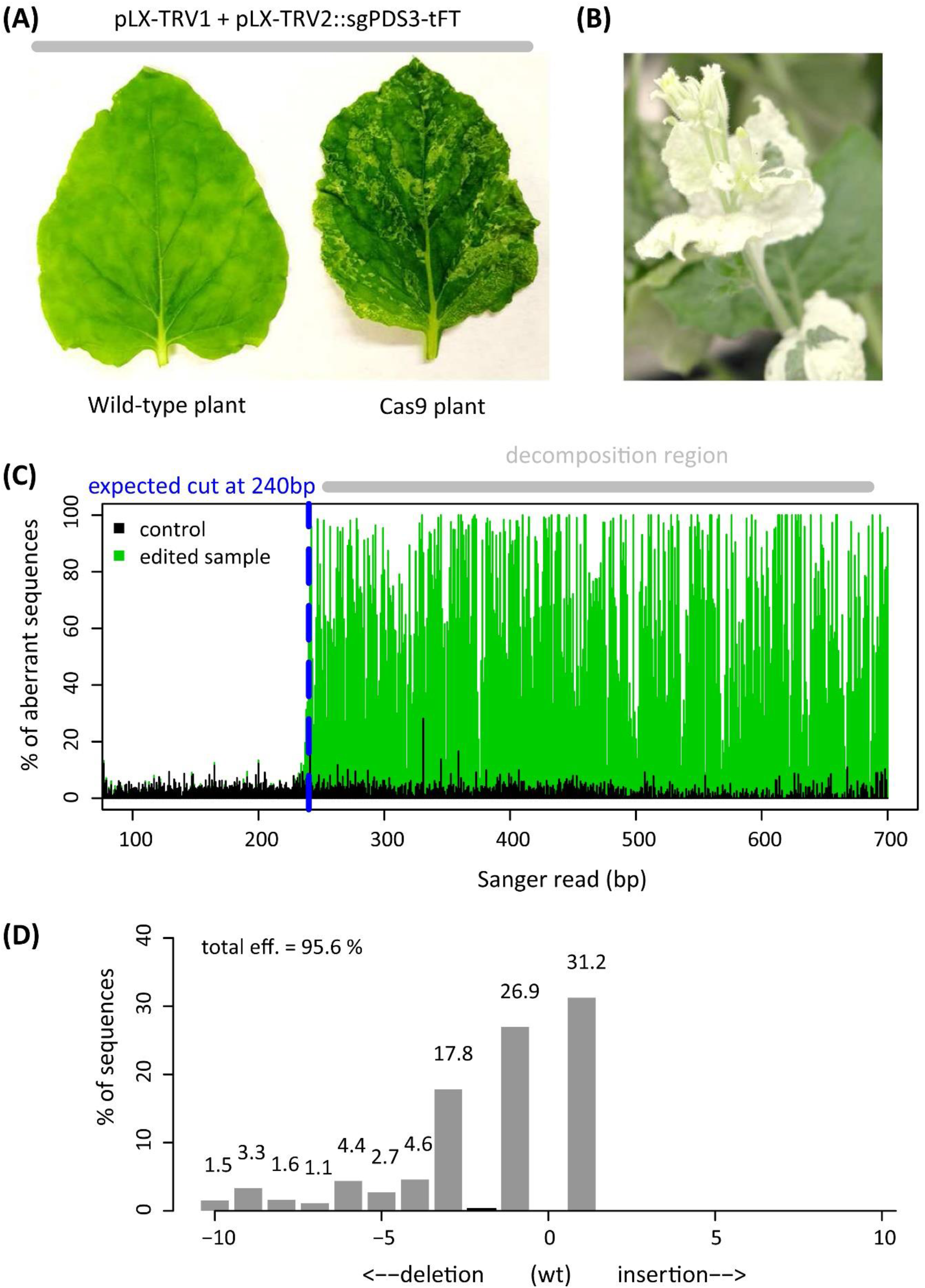
Genome editing of somatic cells using JoinTRV. Plants were agroinoculated with an *Agrobacterium* strain that simultaneously hosts pLX-TRV1 and pLX-TRV2::sgPDS3-tFT, a pLX-TRV2 derivative with a sgRNA targeting two *N. benthamiana* PDS homeologs. **(A)** Phenotypes of agroinoculated wild-type and Cas9 plants (∼1 month post inoculation). **(B)** Severe photobleaching phenotype in Cas9 mature plants (∼2.5 months post inoculation). **(C)** Chromatogram deconvolution of edited and control samples from agroinoculated Cas9 plants; note the high % of aberrant sequence downstream the predicted Cas9 cleavage site (blue dashed line). **(D)** Total editing efficiency and indel distribution at one *PDS* locus of an edited sample (21 days post inoculation). ***Pause point:*** After collection, samples can be stored at -80°C indefinitely.
  b. Purify genomic DNA as described in step 18.b, except for omitting substep 18.b.iii. ***Note:*** Use for downstream applications DNA samples with a ratio of absorbance at 260 nm and 280 nm (260/280) of ∼1.8, and 260/230 ratio in the range of 2.0-2.2. ***Alternative:*** Other procedures and commercial kits for plant genomic DNA purification can also be used. ***Pause point:*** Purified genomic DNA samples can be stored for several weeks at -20°C.
  c. Subject genomic DNA aliquots to PCR amplification with Phusion™ high-fidelity DNA polymerase (Thermo Fisher Scientific) using gene-specific primers.

Phusion™ PCR master mix
| Reagent | Final concentration | Amount |
| --- | --- | --- |
| 5× Phusion™ HF buffer <sup>a</sup> | 1× | 4 µL |
| dNTPs (10 mM each) | 200 µM of each | 0.4 µL |
| Forward primer (10 µM) | 0.5 µM | 1 µL |
| Reverse primer (10 µM) | 0.5 µM | 1 µL |
| Phusion™ high-fidelity DNA polymerase (2U/µL) | 0.02 U/µL | 0.2 µL |
| Genomic DNA | — | ~50 ng |
| Ultrapure water | — | Up to 20 µL |
| **Total** | — | **20 µL** |
| <sup>a</sup> Supplied with the enzyme |  |  |

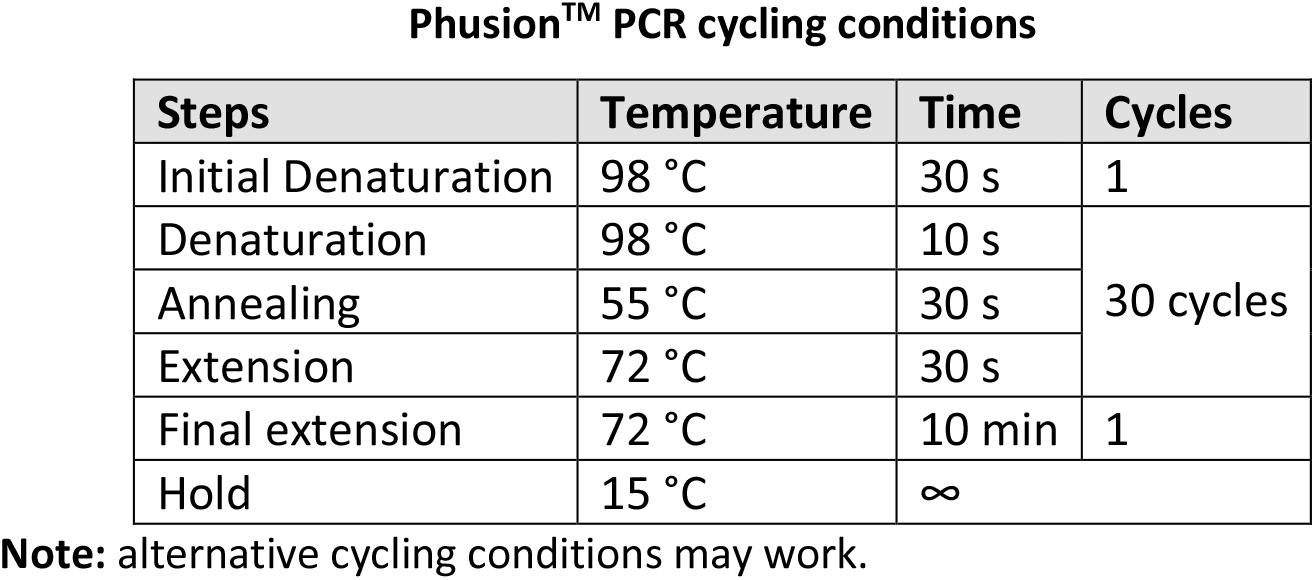 ***Pause point:*** Reactions can be stored for several days at -20°C.
  e. Separate PCR products by electrophoresis in 1% agarose gels in 1× TAE buffer. ***Note:*** A 738-bp fragment of Niben101Scf14708g00023.1 is amplified using the primer pair D4444/D4445.
  f. Purify PCR products from gel bands using the Zymoclean Gel DNA recovery kit (Zymo Research) as per manufacturer’s instructions. ***Alternative:*** Alternative PCR clean-up methods compatible with Sanger sequencing may be used. ***Pause point:*** Purified PCR products can be stored for several days at -20°C.
  g. Subject the purified PCR products to Sanger sequencing. ***CRITICAL:*** Ensure the good quality of sequencing reads by confirming that the electropherograms show high-intensity and well resolved base peaks with no appreciable background noise.
  h. Analyze the presence of targeted modifications within the sequencing reads using the Inference of CRISPR Edits (ICE) (Figure 4C).
    i. Open the online ICE tool (https://ice.synthego.com). ***Alternative:*** Tracking of Indels by Decomposition (TIDE) online tool (https://tide.nki.nl) can also be used for analysis of genome editing; batch analysis may be used (http://shinyapps.datacurators.nl/tide-batch/).
    ii. Paste in “Guide Sequences” the protospacer(s) selected in step 1.c and cloned in the viral vector(s). ***Note:*** Use TTGGTAGTAGCGACTCCATG as the guide target sequence in the analysis of samples obtained using sgPDS3-tFT.
    iii. Upload a Sanger sequencing file from an uninoculated or mock-inoculated plant sample as a reference.
    iv. Upload Sanger sequencing files from the inoculated Cas9 plant samples and retrieve total editing efficiency and indel distribution results. ***Note:*** Samples from infected plants are somatic mosaics of the wild-type genomic sequence and multiple edited alleles (Figure 4D).

### Recovery of mutant progeny

#### Timing: 2-5 months

In this section, progeny of inoculated plants is screened to identify heritable genome editing events.

20. Recovery of mutant progeny (Figure 5).
  a. Collect seeds from inoculated plants. ***CRITICAL:*** Collect seeds from capsules from the top of the parent plant since high fraction of progeny with mutations was reported in capsules distal to the point of infiltration^11^.

**Figure 5.**
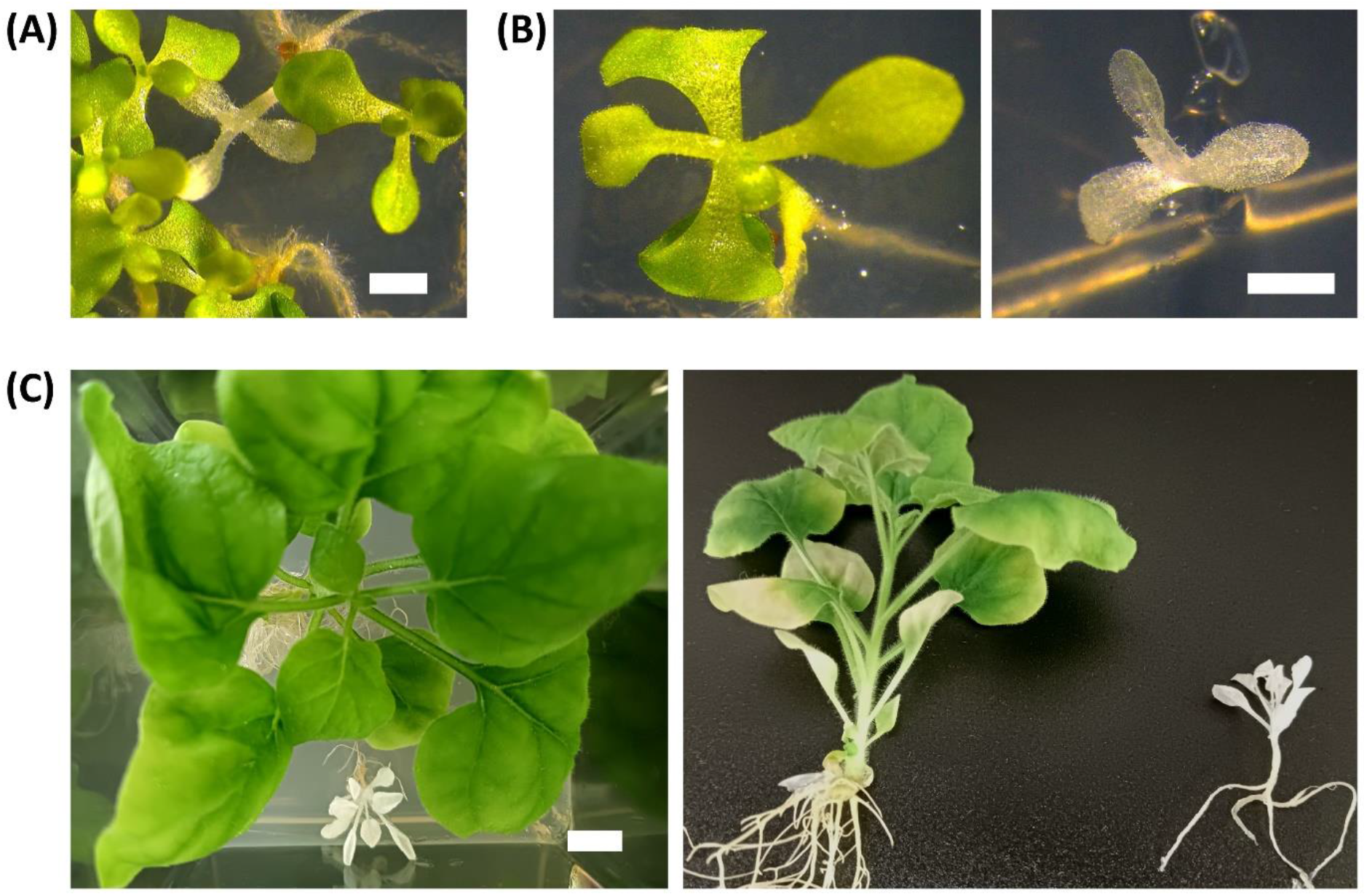
Recovery of mutant progeny using JoinTRV. Cas9 plants were co-inoculated with pLX-TRV1 and pLX-TRV2::sgPDS3-tFT, seeds were collected and the progeny was analyzed. **(A)** Phenotypes of the obtained seedlings at two weeks after sowing; scale bar = 2 mm. **(B)** Details of green and albino seedlings; scale bar = 2 mm. **(C)** Green and albino plantlets at 1.5 months after sowing; scale bar = 1 mm. ***Note:*** On average, 3-4 months are required from plant infection to seed harvest.
  b. Sow seeds on soil and inspect every day for appearance of seedlings with mutant phenotypes. ***CRITICAL:*** Plants with genetic knock-out or chemical inhibition of *PDS* do not survive when grown in soil^36–38^. Sow seeds on ½ MS medium agar plates supplemented with 3% sucrose to facilitate identification of progeny with homozygous loss-of-function mutants of *PDS*.
  c. Collect samples from seedlings and proceed with DNA analysis as described in step 18. ***Note:*** ICE-based indel detection can be used to call genotypes^11^. In progeny analysis, score mutations at each *PDS* homeolog using the following parameters: <35% indels, wild type; 35%–80% indels, heterozygous mutation; >80% indels, homozygous/biallelic mutation (Figure 6).

## Expected outcomes

The procedures described in this protocol enable the assembly of recombinant viral vectors based on binary T-DNA vectors and suitable for *Agrobacterium*-mediated delivery to plants. It is expected that the generated PVX and TRV vectors will be infective and able to spread systemically through the plant, leading to fast (14 dpi) and efficient (up to ≈95% indels) CRISPR-Cas-mediated genome editing in somatic cells^1,2^. This approach conforms an ideal screening tool to rapidly assess sgRNA design at phenotypic and/or genetic levels (Figure 4, Video 1). Finally, targeted modifications are successfully inherited by the progeny of infected plants (Figure 5 and 6).

**Figure 6.**
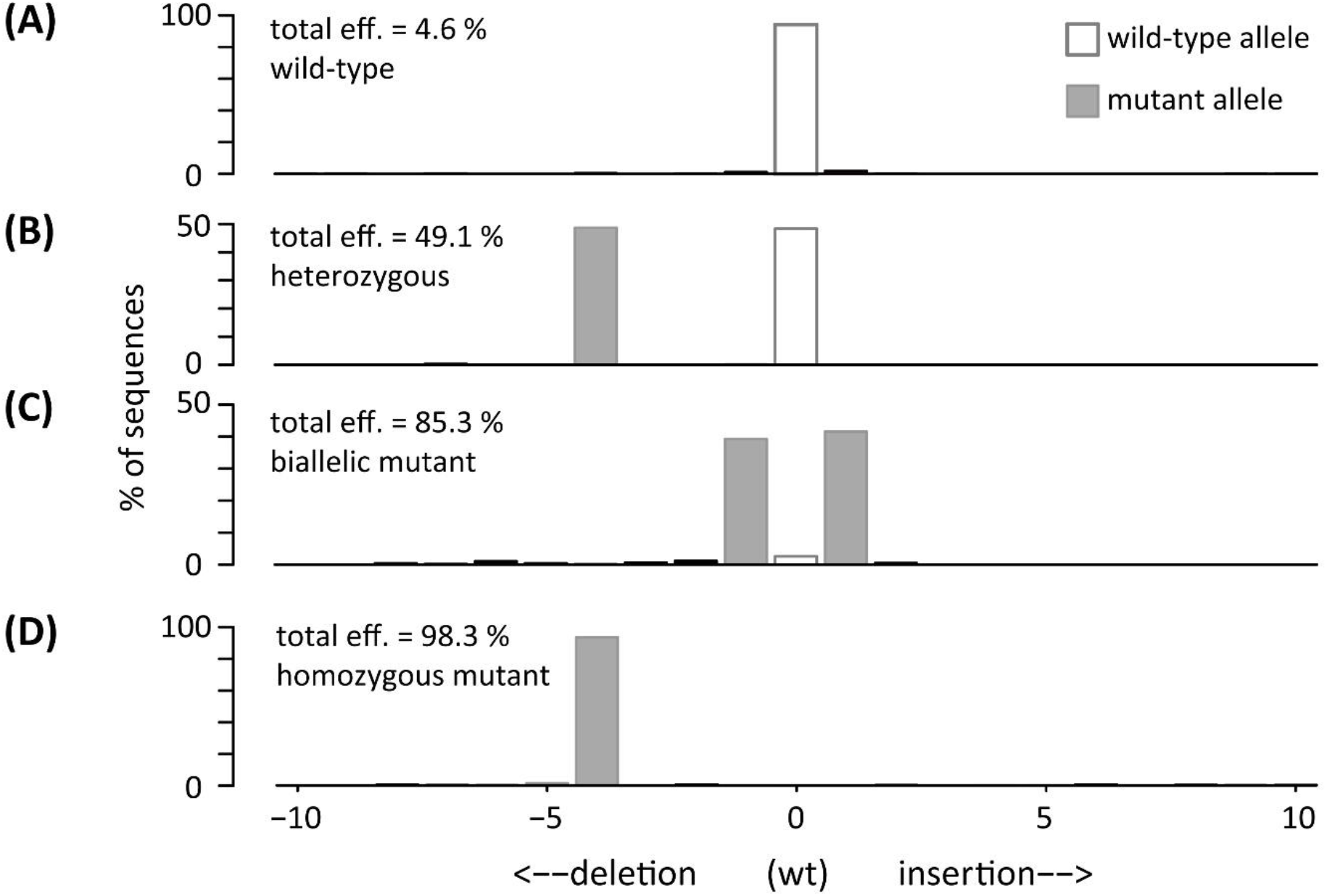
Genotyping of progeny of a *N. benthamiana* Cas9 plant inoculated with JoinTRV. Total editing efficiency and indel distribution at one *PDS* locus of individual progeny lines. Results of representative genotypes are shown: wild-type line (**A**), line with a wild-type allele and a mutant one (**B**), line with two mutant alleles (**C**), homozygous mutant line (**D**).

Besides the reported mutagenesis of targeted genomic loci via double-stranded DNA break induction, the described viral vectors can be used to deliver sgRNA constructs that guide Cas9 fusion proteins for a variety of applications including base editing, transcriptional and metabolic reprogramming^28,39,40^. Furthermore, the described viral vector pLX-PVX and the pLX-TRV1/pLX-TRV2 pair can be used for additional applications such as virus-mediated gene silencing (VIGS) or recombinant protein overexpression^1,2^.

## Limitations

PVX and TRV possess a wide host range specially in the Solanaceae family^41,42^; however, the viral vectors assembled according to this protocol have only been tested in *N. benthamiana*. To apply this protocol to other plant species, generation of a transgenic line stably expressing the *S. pyogenes* Cas9 nuclease is required. A variability in virus infectivity can be observed when inoculating other plant species that may need an optimization of the protocol for each case.

Gene knockout may cause severe growth defects and lethality. Analysis of genes whose loss-of-function phenotypes are lethal may require condition optimization and special selection scheme. Progeny plants recovered from editing assays are transgenic since they include the Cas9 expression cassette.

Prolonged, high levels of exposure to CRISPR–Cas9 delivered using viral vectors have revealed no detectable off-target mutations^17^; however, this aspect should be evaluated on case-by-case basis.

## Troubleshooting

### Problem 1

High background of empty vector colonies precludes identification of positive clones (steps 7-8); no vector is recovered with the correct digestion profile (step 9).

### Potential solutions

Get familiar with procedures for plasmid construct generation and specifically with principles of in vitro homology-based^31^.

Ensure that the cloning strategy and the used materials (i.e. primers, synthetic DNA fragments…) are correct.

Screen more colonies.

Work in sterile conditions and avoid plasmid or bacterial contaminations.

### Problem 2

Low plasmid DNA yield from *E. coli* cultures.

### Potential solutions

Binary vectors used herein include a medium copy origin by design to enhance stability^23^, higher culture volumes (step 8) should be used to reach yields obtained using with high-copy plasmids. The DNA plasmid amount recovered from a single miniprep is nonetheless sufficient for the procedures detailed in this protocol, that is restriction enzyme digestion verification (step 9), clone verification by sequencing (step 10), and transformation of *Agrobacterium* cells (step 11).

### Problem 3

Slow growth rates of bacteria harboring binary vectors with viral vectors.

### Potential solution

For most applications, extended bacterial culturing time may solve the problem.

### Problem 4

Sequence alteration and degeneration of viral vectors after propagation in bacteria.

### Potential solution

Start experiments using a backup plasmid stock.

Bacterial cells harboring binary vectors with full-length viral vectors may display low growth rates. Avoid contaminations with bacterial strains with fast growth rates. If a mixture of small and large colonies appears during agar plate selection pick small colonies for subsequent analysis.

Reduce of *E. coli* culturing temperatures from 37°C to 25-30°C to improve recovery and propagation of full-length viral vectors^43^.

### Problem 5

*N. benthamiana* leaves are hard to infiltrate (step 16).

### Potential solution

Create a small nick with a needle or blade in the epidermis on the adaxial side of the leaf. Make sure not to scratch so hard as to pierce the leaf through, otherwise the *A. tumefaciens* inoculum will pass through the puncture instead of penetrating into leaf tissue.

### Problem 6

Analysis of virus infectivity is inconsistent between plants infected with the same viral vector.

### Potential solution

Use freshly prepared RNA/DNA extraction buffer (see Materials) when performing RNA extractions to prevent false positive results due to contaminations.

Carefully run RT-PCR assays to prevent false positive results due to contaminations with plasmid DNA of the assembled cloned.

### Problem 7

No somatic editing is detected.

### Potential solution

Ensure to use Cas9-transgenic plants.

Ensure that the sgRNA design and the used materials are correct.

Test protospacer sequences designed at difference regions of the target genes.

### Problem 8

PCR products of target genomic regions show poor quality electropherograms (i.e. short reads, multiple peaks) in control, unedited samples.

### Potential solution

*N. benthamiana* is an allotetraploid. For genome editing analysis of specific homeologs ensure to use primer pairs designed and validated by dedicated tools, e.g. BLAST (https://solgenomics.net/tools/blast/) and In Silico PCR (https://solgenomics.net/tools/in_silico_pcr). Resolve PCR products in a 2% agarose gel before to proceed with the gel purification (step 19.e). Replace the gel purification (step 19.f) with alternative PCR clean-up methods compatible with Sanger sequencing.

If the problem persists, clone PCR amplicons into suitable plasmid vectors and perform Sanger sequencing of DNA plasmids from selected colonies.

### Problem 9

No mutant progeny is recovered.

### Potential solution

Somatic editing phenotypes increase in intensity as the plants mature, extending the assay duration and seed collection from late flowers may be required to recover mutant progeny.

Growth conditions of the parental and progeny plants may require empirical optimization to ensure high levels of hereditable editing.

Gene knockout may cause severe growth defects and lethality, which would require large screen efforts or *ad hoc* rescue strategies to identify mutant progeny.

## Resource availability

### Lead contact

Further information and requests for resources and reagents should be directed to and will be fulfilled by the lead contact, José-Antonio Daròs.

### Materials availability

pLX-TRV1 and pLX-TRV2 are available at Addgene with product number 180515 (https://www.addgene.org/180515/) and 180516 (https://www.addgene.org/180516/), respectively.

### Data and code availability

This protocol does not disclose new data or code.

## Acknowledgments

This work was supported by grant PID2020-114691RB-I00 from Ministerio de Ciencia e Innovación (Spain) through the Agencia Estatal de Investigación, and co-financed European Regional Development Fund. M.U. was the recipient of a fellowship (FPU17/05503) from Ministerio de Ciencia e Innovación (Spain). F.P. is financed by a Juan de la Cierva Incorporación contract (IJC2019-039970-I) from Ministerio de Ciencia e Innovación (Spain).

## Author contributions

All authors participated in work conception and design. M.U., F.P. and J.A.D. wrote the manuscript with input from the rest of the authors.

## Figure and video legends

**Video 1**. CRISPR/Cas9 genome editing of plant somatic cells using JoinTRV. Time-course appearance of photobleaching mosaics in *N. benthamiana* Cas9 plants co-inoculated with pLX-TRV1 and pLX-TRV2::sgPDS3-tFT (right). Wild-type plants inoculated with the same vectors are shown as controls (left).

