## Supporting Information for "Heritable CRISPR-Cas9 editing of plant genomes using RNA virus vectors"

### Supplemental sequence 1. pLX-PVX

**>KY825137.1 Cloning vector pLX-B2, complete sequence**

```
CCATCATCAGTTCGGTGGTCTTCCGACGAACAATAAGGCCGCAAATCGCGGCCTTTTTTATTGATAACAAAACCGGCTCAGTTCTGCGTAGAAAC
CAACATGCAAGCTCCACCGGGTGCAAAGCGGCAGCGGCGGCAGGATATATTCAATTGTAAATGGCTTCATGTCCGGGAAATCTACTGGTGGCAGG
ATATATTGTGGTGTAAACAATGGAGAAAAAGATTAATTAAGTACTAGTAGCCTGGTCGAAACCGTCTCAGGAGAGAGACCAAAAGCAAAAACCCG
CCGAAGCGGGTTACTAGCGCCATTCGCCATTCAGAGAGCGGAGCTGCTGCGACGGACGATCGGTACGCGCCTCTTCGCTATTACGCCAACTGGCG
AAAGGTGGATGTGCTGCAAGGCGATTAAGTTGGGTAACGCCAGGGTTTTCCCAGTCACGACGTTGTAGTACCACGGCAAGGCTATCTGTAATCAT
TGTTGTACTCCGGTTAGGACGGATTGGGAACTGGCTAACTCAAAATCCACACATTATACGAGCCGGAAGCATAAAGTGTAAAGCCTGGGGTGCCT
AATGAGTGAGGTCTCTCGCTGTCAGGAGACGGGACAAGGATGCGAGCTAGCCTGCAGGAATTGTTGATTTTGTGATGACTGATGGCAGGATATAT
GCGGTTGTAATTCATTTTTATTGTCTAAATTTCTGTATTTGTTTGTTTGTTCGGTTGTAAATTTTTTTGGAAGAACAAGAAAAGAAAAAACACCC
GTTAGGGTGTTTTTAGTTAGTGTGGCGCGCCGACTTGCGACATGCGGTCCTTTGCAATCAACTATTAGAAAAATTCATCCAGCATCAGATGAAAT
TGCAGTTTGTTCATATCCGGATTATCAATGCCATATTTCTGAAACAGACGTTTTTGCAGGCTCGGGCTAAATTCGCCCAGGCAGTTCCACAGAAT
GGCCAGATCCTGATAACGATCCGCAATGCCCACACGGCCCACATCAATGCAGCCAATCAGTTTGCCTTCATCGAAAATCAGGTTATCCAGGCTAA
AATCGCCGTGGGTCACCACGCTATCCGGGCTAAACGGCAGCAGTTTATGCATTTCTTTCCACACCTGTTCCACCGGCCAGCCGTTACGTTCATCA
TCAAAATCGCTCGCATCCACCAGGCCGTTGTTCATACGGCTCTGCGCCTGGGCCAGACGAAACACACGATCGCTGTTAAACGGGCAGTTGCACAC
CGGAATGCTATGCAGACGACGCAGAAACACGGCCAGCGCATCCACAATGTTTTCGCCGCTATCCGGATATTCTTCCAGCACCTGAAACGCGGTTT
TGCCCGGAATCGCGGTGGTCAGCAGCCACGCATCATCCGGGGTGCGAATAAAATGTTTAATGGTCGGCAGCGGCATAAATTCGGTCAGCCAGTTC
AGACGCACCATTTCATCGGTCACATCGTTCGCCACGCTGCCTTTGCCATGTTTCAGAAACAGTTCCGGCGCATCCGGTTTGCCATACAGACGATA
AATGGTCGCGCCGCTCTGACCCACGTTATCACGCGCCCATTTATAGCCATACAGATCCGCATCCATGTTGCTGTTCAGACGCGGACGGCTACAGC
TCGTTTCACGCTGAATATGGCTCATAACACCCCTTGTATTACTGTTTATGTAAGCAGACAGTTTTATTGTTCATGATGATATATTTTTATCTTGT
GCAATGTAACATCAGAGATTTTGAGACACAAGATCGGATTGGCGGTTATGCGGTTCTACCGGCGCGGCAGCGTTACCCGTGTCGGCGGCTCCAAC
GGCTCGCCATCGTCCAGAAAACACGGCTCATCGGGCATCGGCAGGCGCTGCTGCCCGCGCCGTTCCCATTCCTCCGTTTCGGTCAAGGCTGGCAG
GTCTGGTTCCATGCCCGGAATGCCGGGCTGGCTGGGCGGCTCCTCGCCGGGGCCGGTCGGTAGTTGCTGCTCGCCCGGATACAGGGTCGGGATGC
GGCGCAGGTCGCCATGCCCCAACAGCGATTCGTCCTGGTCGTCGTGATCAACCACCACGGCGGCACTGAACACCGACAGGCGCAACTGGTCGCGG
GGCTGGCCCCACGCCACGCGGTCATTGACCACGTAGGCCGACACGGTGCCGGGGCCGTTGAGCTTCACGACGGAGATCCAGCGCTCGGCCACCAA
GTCCTTGACTGCGTATTGGACCGTCCGCAAAGAACGTCCGATGAGCTTGGAAAGTGTCTTCTGGCTGACCACCACGGCGTTCTGGTGGCCCATCT
GCGCCACGAGGTGATGCAGCAGCATTGCCGCCGTGGGTTTCCTCGCAATAAGCCCGGCCCACGCCTCATGCGCTTTGCGTTCCGTTTGCACCCAG
TGACCGGGCTTGTTCTTGGCTTGAATGCCGATTTCTCTGGACTGCGTGGCCATGCTTATCTCCATGCGGTAGGGGTGCCGCACGGTTGCGGCACC
ATGCGCAATCAGCTGCAACTTTTCGGCAGCGCGACAACAATTATGCGTTGCGTAAAAGTGGCAGTCAATTACAGATTTTCTTTAACCTACGCAAT
GAGCTATTGCGGGGGGTGCCGCAATGAGCTGTTGCGTACCCCCCTTTTTTAAGTTGTTGATTTTTAAGTCTTTCGCATTTCGCCCTATATCTAGT
TCTTTGGTGCCCAAAGAAGGGCACCCCTGCGGGGTTCCCCCACGCCTTCGGCGCGGCTCCCCCTCCGGCAAAAAGTGGCCCCTCCGGGGCTTGTT
GATCGACTGCGCGGCCTTCGGCCTTGCCCAAGGTGGCGCTGCCCCCTTGGAACCCCCGCACTCGCCGCCGTGAGGCTCGGGGGGCAGGCGGGCGG
GCTTCGCCCTTCGACTGCCCCCACTCGCATAGGCTTGGGTCGTTCCAGGCGCGTCAAGGCCAAGCCGCTGCGCGGTCGCTGCGCGAGCCTTGACC
CGCCTTCCACTTGGTGTCCAACCGGCAAGCGAAGCGCGCAGGCCGCAGGCCGGAGGCTTTTCCCCAGAGAAAATTAAAAAAATTGATGGGGCAAG
GCCGCAGGCCGCGCAGTTGGAGCCGGTGGGTATGTGGTCGAAGGCTGGGTAGCCGGTGGGCAATCCCTGTGGTCAAGCTCGTGGGCAGGCGCAGC
CTGTCCATCAGCTTGTCCAGCAGGGTTGTCCACGGGCCGAGCGAAGCGAGCCAGCCGGTGGCCGCTCGCGGCCATCGTCCACATATCCACGGGCT
GGCAAGGGAGCGCAGCGACCGCGCAGGGCGAAGCCCGGAGAGCAAGCCCGTAGGGGG
```

**>MT799816.1 Potato virus X isolate PVX.a, complete genome**

```
GAAAACTAAACCATACACCACCAACACAACCAAACCCACCACGCCCAATTGTTACACACCCGCTTGAAAAAGAAAGTTTAACAAATGGCCAAGGT
GCGCGAGGTTTACCAATCTTTTACAGACTCCACCACAAAAACTCTCATCCAAGATGAGGCTTATAGAAACATTCGCCCCATCATGGAAAAACACA
AACTAGCTAACCCTTACGCTCAAACGGTTGAAGCGGCTAATGATCTAGAGGGGTTCGGCATAGCCACCAATCCCTATAGCATTGAATTGCATACA
CATGCAGCCGCTAAGACCATAGAGAATAAACTTCTAGAGGTGCTTGGTTCCATCCTACCACAAGAACCTGTTACATTTATGTTTCTTAAACCCAG
AAAGCTAAACTACATGAGAAGAAACCCGCGGATCAAGGACATTTTCCAAAATGTTGCCATTGAACCAAGAGACGTAGCCAGGTACCCCAAGGAAA
CAATAATTGACAAACTCACAGAGATCACAACGGAAACAGCATACATTAGTGACACTCTGCACTTCTTGGATCCGAGCTACATAGTGGAGACATTC
CAAAACTGCCCAAAATTGCAAACATTGTATGCGACCTTAGTTCTCCCCGTTGAGGCAGCCTTTAAAATGGAAAGCACTCACCCGAACATATACAG
CCTCAAATACTTCGGAGATGGTTTCCAGTATATACCAGGCAACCATGGTGGCGGGGCATACCATCATGAATTCGCTCATCTACAATGGCTCAAAG
TGGGAAAGATCAAGTGGAGGGACCCCAAGGATAGCTTTCTCGGACATCTCAATTACACGACTGAGCAGGTTGAGATGCACACAGTGACAGTACAG
TTGCAGGAATCGTTCGCGGCAAACCACTTGTACTGCATCAGGAGAGGAGACTTGCTCACACCGGAGGTGCGCACTTTCGGCCAACCTGACAGGTA
CGTGATTCCACCACAGATCTTCCTCCCAAAAGTTCACAACTGCAAGAAGCCGATTCTCAAGAAAACTATGATGCAGCTCTTCTTGTATGTTAGGA
CAGTCAAGGTCGCAAAAAATTGTGACATTTTTGCCAAAGTCAGACAATTAATTAAATCATCTGACTTGGACAAATACTCTGCTGTGGAACTGGTT
TACTTAGTAAGCTACATGGAGTTCCTTGCCGATTTACAAGCTACCACCTGCTTCTCAGACACACTTTCTGGTGGCTTGCTAACAAAGACCCTTGC
ACCGGTGAGGGCTTGGATACAAGAGAAAAAGATGCAGCTGTTTGGTCTTGAGGACTACGCGAAGTTAGTCAAAGCAGTTGATTTCCACCCGGTGG
ATTTTTCTTTCAAAGTGGAAACTTGGGACTTCAGATTCCACCCCTTGCAAGCGTGGAAAGCCTTCCGACCAAGGGAAGTGTCGGATGTAGAGGAA
ATGGAAAGTTTGTTCTCAGATGGGGACTTGCTTGATTGCTTCACAAGAATGCCAGCTTATGCGGTAAACGCAGAGGAAGATTTAGCTGCAATCAG
GAAAACGCCCGAGATGGATGTCGGTCAAGAAGTTAAAGAACCTGCAGGAGACAGAAATCAATACTCAAACCCTGCAGAAACTTTCCTCAACAAGC
TCCACAGGAAACACAGTAGGGAGGTGAAACACCAGGCCGCAAAGAAAGCTAAACGCCTAGCTGAAATCCAGGAGTCAATGAGAGCTGAAGGTGAT
GCCGAACCAAATGAAATAAGCGGGGGCATGGGGGCAATACCCAGCAACGCCGAACTTCCTGGCACGAATGATGCCAGACAAGAACTCACACTCCC
AACCACTAAACCTGTCCCTGCAAGGTGGGAAGATGCTTCATTCACAGATTCTAGTGTGGAAGAGGAGCAGGTTAAACCCCTTGGAAAAGAAACCG
TTGAAACAGCGACGCAACAAGTCATCGAAGGACTTCCTTGGAAACACTGGATTCCTCAATTAAATGCTGTTGGATTCAAGGCGCTGGAAATTCAG
AGGGATAGGAGTGGAACAATGATCATGCCCATCACAGAAATGGTCTCCGGGCTGGAAAAAGAGGACTTCCCTGAAGGAACTCCAAAAGAGTTGGC
ACGAGAATTGTTCGCTATGAACAGAAGCCCTGCCACCATCCCTTTGGACCTGCTTAGAGCCAGAGACTACGGCAGTGATGTAAAGAACAAGAGAA
TTGGTGCCATCACAAAGACACAGGCAACGAGTTGGGGCGAATACTTGACAGGAAAGATAGAAAGCTTGACTGAGAGGAAAGTTGCGACTTGTGTC
ATTCATGGAGCTGGAGGTTCTGGAAAAAGTCATGCCATCCAGAAGGCATTGAGAGAAATTGGCAAGGGCTCGGACATCACTGTAGTCCTGCCGAC
CAATGAACTGCGGCTAGATTGGAGTAAGAAAGTGCCTAACACTGAGCCCTATATGTTCAAGACCTATGAAAAGGCGTTAATTGGGGGAACAGGCA
GCATAGTCATCTTTGACGATTACTCAAAACTTCCTCCCGGTTACATAGAAGCCTTAGTCTGTTTCTACTCTAAAATCAAGCTAATCATTCTAACA
GGAGATAGCAGACAAAGCGTCTACCATGAAACTGCTGAGGACGCCTCCATCAGGCATTTGGGACCAGCAACAGAGTACTTCTCAAAATACTGCCG
ATACTATCTCAATGCCACACACCGCAACAAGAAAGATCTTGCGAACATGCTTGGTGTCTACAGTGAGAGAACGGGAGTCACCGAAATCAGCATGA
```

GCGCCGAGTTCTTAGAAGGAATCCCAACTTTGGTACCCTCGGATGAGAAGAGAAAGCTGTACATGGGCACCGGGAGGAATGACACGTTCACATAC
GCTGGATGCCAGGGGCTAACTAAGCCGAAGGTACAAATAGTGTTGGACCACAACACCCAAGTGTGTAGCGCGAATGTGATGTACACGGCACTTTC
TAGAGCCACCGATAGGATTCACTTCGTGAACACAAGTGCAAATTCCTCGGCCTTCTGGGAAAAGTTGGACAGCACCCCTTACCTCAAGACTTTCC
TATCAGTGGTGAGAGAACAAGCACTCAGGGAGTACGAGCCGGCAGAGGCAGAGCCAATTCAAGAGCCTGAGCCCCAGACACACATGTGTGTCGAG
AATGAGGAGTCCGTGCTAGAAGAGTACAAAGAGGAACTCTTGGAAAAGTTTGACAGAGAGATCCACTCTGAATCCCATGGTCATTCAAACTGTGT
CCAAACTGAAGACACAACCATTCAGTTGTTTTCGCATCAACAAGCAAAAGATGAGACCCTCCTCTGGGCGACTATAGATGCGCGGCTCAAGACCA
GCAATCAAGAGACAAACTTCCGAGAATTCCTGAGCAAGAAGGACATTGGGGACGTTCTGTTTTTAAACTACCAAAAAGCTATGGGTTTACCCAAA
GAGCGTATTCCTTTTTCCCAAGAGGTCTGGGAAGCTTGTGCCCACGAAGTACAAAGCAAGTACCTCAGCAAGTCAAAGTGCAACTTGATCAATGG
GACTGTGAGACAGAGCCCAGACTTCGATGAAAATAAGATTATGGTATTCCTCAAGTCGCAGTGGGTCACAAAGGTGGAAAAACTAGGTCTACCCA
AGATTAAGCCAGGTCAAACCATAGCAGCCTTTTACCAGCAGACTGTGATGCTTTTTGGAACTATGGCTAGGTACATGCGATGGTTCAGACAGGCT
TTCCAGCCAAAAGAAGTCTTCATAAACTGTGAGACCACGCCAGATGACATGTCTGCATGGGCCTTGAACAACTGGAATTTCAGCAGACCTAGCTT
GGCTAATGACTACACAGCTTTCGACCAGTCTCAGGATGGAGCCATGTTGCAATTTGAGGTGCTCAAAGCCAAACACCACTGCATACCAGAGGAAA
TCATTCAGGCATACATAGATATTAAGACTAATGCACAGATTTTCCTAGGCACGTTATCAATTATGCGCCTGACTGGTGAAGGTCCCACTTTTGAT
GCAAACACTGAGTGCAACATAGCTTACACCCATACAAAGTTTGACATCCCAGCCGGAACTGCTCAAGTTTATGCAGGAGACGACTCCGCACTGGA
CTGTGTTCCAGAAGTGAAGCATAGTTTCCACAGGCTTGAGGACAAATTACTCCTAAAGTCAAAGCCTGTAATCACGCAGCAAAAGAAGGGCAGTT
GGCCTGAGTTTTGTGGTTGGCTGATCACACCAAAAGGGGTGATGAAAGACCCAATTAAGCTCCATGTTAGCTTAAAATTGGCTGAAGCTAAGGGT
GAACTCAAGAAATGTCAAGATTCCTATGAAATTGATCTGAGTTATGCCTATGACCACAAGGACTCTCTGCATGACTTGTTCGATGAGAAACAGTG
TCAGGCACACACACTCACTTGCAGAACACTAATCAAGTCAGGGAGAGGCACTGTCTCACTTTCCCGCCTCAGAAACTTTCTTTAACCGTTAAGTT
ACCTTAGAGATTTGAATAAGATGGATATTCTCATCAGTAGTTTGAAAAGTTTAGGTTATTCTAGGACTTCCAAATCTTTAGATTCAGGACCTTTG
GTAGTACATGCAGTAGCCGGAGCCGGTAAGTCCACAGCCCTAAGGAAGTTGATCCTCAGACACCCAACATTCACCGTGCATACACTCGGTGTCCC
TGACAAGGTGAGTATCAGAACTAGAGGCATACAGAAGCCAGGACCTATTCCTGAGGGCAACTTCGCAATCCTCGATGAGTATACTTTGGACAACA
CCACAAGGAACTCATACCAGGCACTTTTTGCTGACCCTTATCAGGCACCGGAGTTTAGCCTAGAGCCCCACTTCTACTTGGAAACATCATTTCGA
GTTCCGAGGAAAGTGGCAGATTTGATAGCTGGCTGTGGCTTCGATTTCGAGACCAACTCACCGGAAGAAGGGCACTTAGAGATCACTGGCATATT
CAAAGGGCCCCTACTCGGAAAGGTGATAGCCATTGATGAGGAGTCTGAGACAACACTGTCCAGGCATGGTGTTGAGTTTGTTAAGCCCTGCCAAG
TGACGGGACTTGAGTTCAAAGTAGTCACTATTGTGTCTGCCGCACCAATAGAGGAAATTGGCCAGTCCACAGCTTTCTACAACGCTATCACCAGG
TCAAAGGGATTGACATATGTCCGCGCAGGGCCATAGGCTGACCGCTCCGGTCAATTCTGAAAAAGTGTACATAGTATTAGGTCTATCATTTGCTT
TAGTTTCAATTACCTTTCTGCTTTCTAGAAATAGCTTACCCCACGTCGGTGACAACATTCACAGCTTGCCACACGGAGGAGCTTACAGAGACGGC
ACCAAAGCAATCTTGTACAACTCCCCAAATCTAGGGTCACGAGTGAGTCTACACAACGGAAAGAACGCAGCATTTGCTGCCGTTTTGCTACTGAC
TTTGCTGATCTATGGAAGTAAATACATATCTCAACGCAATCATACTTGTGCTTGTGGTAACAATCATAGCAGTCATTAGCACTTCCTTAGTGAGG
ACTGAACCTTGTGTCATCAAGATTACTGGGGAATCAATCACAGTGTTGGCTTGCAAACTAGATGCAGAAACCATAAGGGCCATTGCCGATCTCAA
GCCACTCTCCGTTGAACGGTTAAGTTTCCATTGATACTCGAAAGATGTCAGCACCAGCTAGCACAACACAGCCCATAGGGTCAACTACCTCAACT
ACCACAAAAACTGCAGGCGCAACTCCTGCCACAGCTTCAGGCCTGTTCACTATCCCGGATGGGGATTTCTTTAGTACAGCCCGTGCCATAGTAGC
CAGCAATGCTGTCGCAACAAATGAGGACCTCAGCAAGATTGAGGCTATTTGGAAGGACATGAAGGTGCCCACAGACACTATGGCACAGGCTGCTT
GGGACTTAGTCAGACACTGTGCTGATGTAGGATCATCCGCTCAAACAGAAATGATAGATACAGGTCCCTATTCCAACGGCATCAGCAGAGCTAGA
CTGGCAGCAGCAATTAAAGAGGTGTGCACACTTAGGCAATTTTGCATGAAGTATGCCCCAGTGGTATGGAACTGGATGTTAACTAACAACAGTCC
ACCTGCTAACTGGCAAGCACAAGGTTTCAAGCCTGAGCACAAATTCGCTGCATTCGACTTCTTCAATGGAGTCACCAACCCAGCTGCCATCATGC
CCAAAGAGGGGCTCATCCGGCCACCGTCTGAAGCTGAAATGAATGCTGCCCAAACTGCTGCCTTTGTGAAGATTACAAAGGCCAGGGCACAATCC
AACGACTTTGCCAGCCTAGATGCAGCTGTCACTCGAGGTCGTATCACTGGAACAACAACCGCTGAGGCTGTTGTCACTCTACCACCACCATAACT
ACGTCTACATAACCGACGCCTACCCCAGTTTCATAGTATTTTCTGGTTTGATTGTATGAATAATATAAAT
